# Dictionary learning for integrative, multimodal, and scalable single-cell analysis

**DOI:** 10.1101/2022.02.24.481684

**Authors:** Yuhan Hao, Tim Stuart, Madeline Kowalski, Saket Choudhary, Paul Hoffman, Austin Hartman, Avi Srivastava, Gesmira Molla, Shaista Madad, Carlos Fernandez-Granda, Rahul Satija

## Abstract

Mapping single-cell sequencing profiles to comprehensive reference datasets represents a powerful alternative to unsupervised analysis. Reference datasets, however, are predominantly constructed from single-cell RNA-seq data, and cannot be used to annotate datasets that do not measure gene expression. Here we introduce ‘bridge integration’, a method to harmonize singlecell datasets across modalities by leveraging a multi-omic dataset as a molecular bridge. Each cell in the multi-omic dataset comprises an element in a ‘dictionary’, which can be used to reconstruct unimodal datasets and transform them into a shared space. We demonstrate that our procedure can accurately harmonize transcriptomic data with independent single cell measurements of chromatin accessibility, histone modifications, DNA methylation, and protein levels. Moreover, we demonstrate how dictionary learning can be combined with sketching techniques to substantially improve computational scalability, and harmonize 8.6 million human immune cell profiles from sequencing and mass cytometry experiments. Our approach aims to broaden the utility of single-cell reference datasets and facilitate comparisons across diverse molecular modalities.

**Availability:** Installation instructions, documentations, and vignettes are available at http://www.satijalab.org/seurat

## Introduction

In the same way that read mapping tools have transformed genome sequence analysis^1–3^, the ability to map new datasets to established references represents an exciting opportunity for the field of single-cell genomics. As an alternative to fully unsupervised clustering, supervised mapping approaches leverage large and well-curated references to interpret and annotate query profiles. This strategy is enabled by the curation and public release of reference datasets, as well as the development of new computational tools, including statistical learning^4–7^ and deep learningbased approaches^8,9^ that have been successfully applied towards this goal.

While powerful, a significant current limitation of existing approaches is their primary focus on single-cell RNA-seq (scRNA-seq) data. Single cell transcriptomics is well-suited for the assembly and annotation of reference datasets, particularly as differentially expressed gene markers can typically be interpreted to help annotate cell clusters. This has led to the development of high-quality, carefully curated, and expertly annotated references, particularly from consortia including the Human Cell Atlas (HCA)^10^, Human Biomolecular Atlas Project (HuBMAP^11^), and the Chan Zuckerberg Biohub^12^. Mapping to these references facilitates data harmonization, standardization of cell ontologies and naming schemes, and comparison of scRNA-seq datasets across experimental conditions and disease states.

A crucial challenge is to extend reference-mapping to additional molecular modalities, including single-cell measurements of chromatin accessibility (e.g. scATAC-seq^13,14^), DNA methylation (scBS-seq^15^), histone modifications (scCUT&Tag^16,17^), and protein levels (CyTOF^18^), each of which measures a different set of features than scRNA-seq. The lack of transcriptome-wide measurements creates challenges for unsupervised annotation. Ideally, datasets from different modalities could be mapped onto scRNA-seq references, ensuring that established cell labels and ontologies would be preserved. We and others have proposed methods to map datasets across modalities^19–21^, but these make strict biological assumptions (for example, that accessible chromatin is associated with active transcription) that may not always hold true, particularly when analyzing cellular transitions or developmental trajectories^22^.

Here we introduce ‘bridge integration’, which performs integration of single-cell datasets measuring different modalities by leveraging a separate dataset where both modalities are simultaneously measured as a molecular ‘bridge’. The multi-omic bridge dataset, which can be generated by a diverse set of technologies^23–32^ (**Fig. 1a**), helps to translate information between disparate measurements, resulting in robust integration without requiring any limiting biological assumptions. We illustrate the broad applicability of our approach, demonstrating its performance across five different molecular modalities, and highlighting specific requirements for the multiomic dataset that can help to guide experimental design.

**Figure 1.**
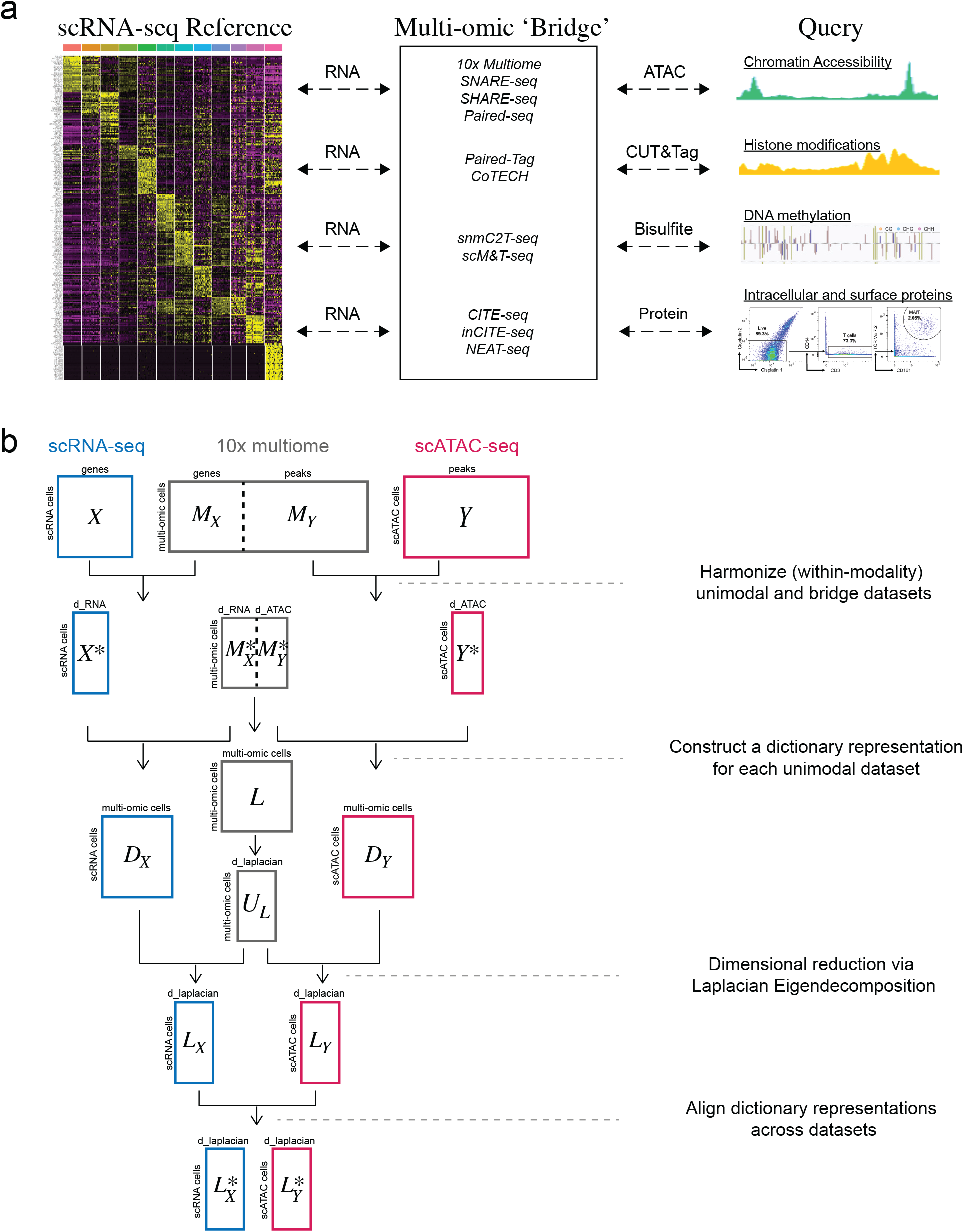
Integrating across modalities with molecular bridges. (**a**) Broad schematic of bridge integration workflow. Two datasets where different modalities are measured (e.g. scRNA-seq and scATAC-seq), can be harmonized via a third dataset where both modalities are simultaneously measured (e.g. 10x multiome). We demonstrate bridge integration using a variety of multi-omic technologies that can be used as bridges, including 10x multiome, Paired-Tag, snmC2T, and CITE-seq, each of which facilitates integration with a different molecular modality. Middle box lists alternative multi-omic technologies that can be used to generate bridge datasets. (**b**) Mathematical schematic of each of the steps in the bridge integration procedure. A full description is provided in the Supplementary Methods. For clarity, the matrix names illustrated in this schematic are the same as the matrix names defined in the Supplementary Methods.

Bridge integration leverages tools from a subfield of representation learning known as ‘dictionary learning’, which are commonly used in image analysis^33^. The goal of dictionary learning is to find a representation of the input data as a weighted linear combination of individual basic elements. We show that dictionary learning has multiple potential applications for single-cell analysis. Our bridge integration procedure is enabled by treating each cell in a multi-omic dataset as elements of a dictionary that can be utilized to reconstruct single-modality datasets. Moreover, we demonstrate how the development of compact dictionaries via dataset sketching can dramatically improve the computational efficiency of large-scale single-cell analysis, and enable rapid integration of dozens of datasets spanning millions of cells.

## Results

We aimed to develop a flexible and robust integration strategy to integrate data from single-cell sequencing experiments where different modalities are measured (‘single-modality datasets’). The fundamental challenge is that different single-modality datasets measure different sets of features. For example, scRNA-seq measures the expression level of individual transcripts, while scATAC-seq or scBS-seq measure DNA accessibility or methylation levels (**Fig. 1a**). Previously proposed methods from our group and others^19–21^ attempt to convert one set of features into another, for example, taking the gene-body sum of ATAC-seq signal (or the inverse of the DNA methylation levels), as a proxy for transcriptional output. While this conversion facilitates downstream integration, it assumes a strict and simplistic biological relationship between modalities that may not hold true, particularly in developing or transitioning systems.

### Utilizing multi-omic dictionaries for bridge integration

We reasoned that an alternative approach would be to leverage a multi-omic dataset as a bridge that can help to translate between disparate modalities. To perform this translation, we were inspired by the field of dictionary learning, a form of representation learning that is commonly utilized in image analysis and also genomics^33–36^. The goal of dictionary learning is to represent input data, a noisy image for example, in terms of individual elements. These elements, such as image patches, are called atoms and together comprise a dictionary. Reconstructing an image as a weighted linear combination of these atoms is an effective tool for denoising, and represents a transformation of the image dataset into a dictionary-defined space.

We find that dictionary learning is a powerful tool for enabling cross-modality bridge integration at single-cell resolution. Our key insight is to treat a multi-omic dataset as a dictionary, with each individual cell’s multi-omic profile representing an atom. We learn a ‘dictionary representation’ of each unimodal dataset based on these atoms. This transformation takes datasets in which completely different sets of features were measured and represents them each in a space where the defining features represent the same set of atoms (**Fig. 1b**). Once different modalities can be represented using the same set of features, they can be readily aligned in a final step.

Our bridge integration is illustrated in **Fig. 1b** and described fully in the Supplementary Methods, and we note a few key points below. First, our procedure makes no assumptions about the relationships between modalities, as these are learned automatically from the multi-omic dataset. Second, the key advance we present here is a transformation to project datasets profiling different modalities to be represented by a shared set of features. Once transformed, the final alignment step is compatible with a wide diversity of single-cell integration techniques including Harmony^37^, mnnCorrect^38^, Seurat^19^, Scanorama^39^, or scVI^40^. In this manuscript, we perform this step with an implementation of the mnnCorrect algorithm^38^.

Lastly, we found that when working with sizable bridge datasets, the large number of atoms (single cells in the bridge dataset) created a substantial computational burden. Motivated by a similar problem addressed by Laplacian Eigenmaps^41^, we compute the graph laplacian for the multi-omic dataset, and calculate an eigendecomposition, thereby reducing the dimensionality from the number of atoms to the number of selected eigenvectors (Supplementary Methods). We then utilize these eigenvectors to transform the learned dictionary representations into the same lower-dimensional space, substantially increasing the efficiency of our bridge integration procedure.

### Mapping scATAC-seq data onto scRNA-seq references

We first demonstrate our bridge integration strategy by performing cross-modality mapping on scATAC-seq and scRNA-seq samples of human bone marrow mononuclear cells (BMMCs). These samples consist of cells representing the full spectrum of hematopoietic differentiation, including hematopoietic stem cells, multi and oligopotent progenitors, and fully differentiated cells. As part of HuBMAP, we have leveraged public datasets to construct a comprehensive scRNA-seq reference (‘Azimuth reference’; 297,627 cells) of human BMMC, carefully annotating 10 progenitor and 25 differentiated cell states (**Fig. 2a**). We aimed to map scATAC-seq ‘query’ datasets of human BMMC^42^ (16,266 whole bone marrow profiles and 9,893 CD34^+^ enriched profiles) to this reference (**Fig. 2b**). We used a 10x Multiome dataset^43^ (32,368 cells paired snRNA-seq + scATAC-seq) that was publicly released as part of NeurIPS 2021, as a bridge.

**Figure 2.**
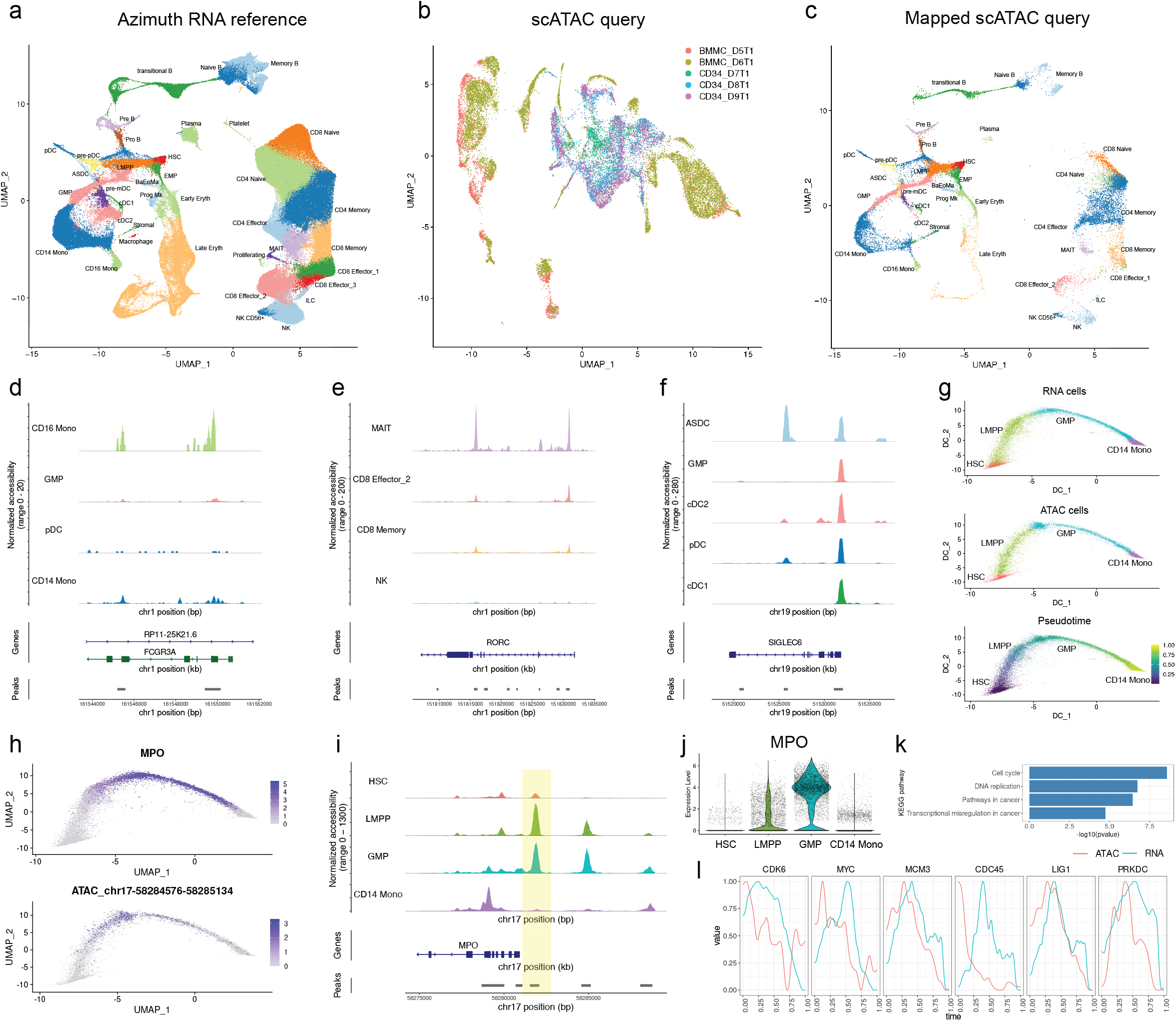
Mapping scATAC-seq data onto scRNA-seq references. (**a**) UMAP visualization of scRNA-seq reference dataset of human bone marrow, representing 297,627 annotated cells. (**b**) UMAP visualization of an scATAC-seq query dataset from (Granja et al, 2019), representing 26,159 profiles spanning five batches, three of which are enriched for CD34 expressing cells. (**c**) After bridge integration, query cells are annotated based on the scRNA-seq defined cell ontology, and can be visualized on the same embedding. (**d-f**) Coverage plots showing chromatin accessibility at selected loci, after grouping query cells by their predicted annotations. In each case, the predicted cell labels agree with the expected accessibility patterns. (**g**) We constructed a differentiation trajectory and pseudotime ordering of cells undergoing myeloid differentiation. The pseudotime ordering encompasses both scRNA-seq and scATAC-seq cells. (**h**) Example locus where we observe a ‘lag’ between the gene expression dynamics for MPO and the accessibility dynamics for an upstream regulatory region (denoted by a yellow box in (**i**)). **(i**) chromatin accessibility at the MPO regulatory locus. The highlighted region becomes accessible at the multipotent LMPP stage. **(j)** MPO becomes highly expressed at the RNA level at the myeloid-committed GMP stage. (**k**) KEGG pathway enrichment for 236 genes where we identified a lag between accessibility and transcriptional dynamics. (**l**) Smoothed chromatin accessibility levels (red) and lagging expression of associated genes (blue) as a function of pseudotime, for 6 cell cycle-associated genes.

Our bridge procedure successfully mapped the scATAC-seq dataset on our Azimuth reference, enabling joint visualization and annotation of scATAC-seq and scRNA-seq data (**Fig. 2c**). Reference-mapping also aligned shared cell populations across multiple samples, mitigating sample-specific batch effects. Query samples representing CD34^+^ BMMC fractions mapped exclusively to the HSC and progenitor components in the reference dataset, demonstrating that bridge integration can robustly handle cases where the query dataset represent a subset of the reference, while whole fractions mapped to all 35 cell states (**Supplementary Fig. 1a**).

Our reference-derived annotations were concordant with the annotations accompanying the query dataset produced by the original authors (**Supplementary Fig. 1b**), but we found that bridge integration annotated additional rare and high-resolution subpopulations. For example, our annotations separated monocytes into CD14^+^ and CD16^+^ fractions, NK cells into CD56^bright^ and CD56^dιm^ subgroups, and cytotoxic T cells into CD8^+^ and mucosal associated invariant T (MAIT) subpopulations (**Fig. 2d,e** and **Supplementary Fig. 1c,d**). While these subdivisions were not identified in the unsupervised scATAC-seq analysis, we confirmed these predictions by observing differential accessibility at canonical loci (i.e. elevated accessibility at the FCGR3A/CD16 gene locus in CD16^+^ monocytes), after grouping by reference-derived annotations. Similarly, bridge integration identified extremely rare groups of innate lymphoid cells (ILC; 0.15%), and recently discovered AXL^+^SIGLEC6^+^ (ASDC) dendritic cells^44,45^ (0.10%) (**Fig. 2f** and **Supplementary Fig. 1e,f**). To our knowledge, these cell populations have not been previously identified in scATAC-seq data. Again, we found that differentially accessible sites, such as an ASDC-specific peak in the SIGLEC6 gene (**Fig. 2f**), fully supported the accuracy of our mapping procedure.

Our reference-mapping procedure not only enables the transfer of discrete annotations, but by projecting datasets from multiple modalities into a common space, allows us to explore how variation in one corresponds to variation in another. For example, after integration, we applied diffusion maps to the harmonized measurements to construct a joint differentiation trajectory spanning multiple progenitor states during myeloid differentiation (**Fig. 2g**). Since this trajectory represents both reference and query cells, we can explore how pseudo-temporal variation in chromatin accessibility correlates with gene expression, even though the two modalities were measured in separate experiments.

Consistent with previous findings, we identified cases where gene expression changes ‘lagged’ behind variation in chromatin accessibility. For example, while myeloperoxidase (MPO) expression is expressed in granulocyte-macrophage progenitors (GMP) and is associated with myeloid fate commitment^46,47^, the regulatory region immediately upstream acquired accessibility in lymphoid-primed multipotent progenitors (LMPP) (**Fig. 2h-j)**. We utilized a cross-correlation-based metric to systematically identify 236 ‘lagging’ loci (Supplementary Methods) across this trajectory. KEGG pathway enrichment analysis revealed a strong enrichment for genes involved in cell cycle and DNA replication (**Fig. 2k**). These loci were characterized by accessible chromatin at the earliest stages of differentiation (hematopoietic stem cells), but there is a delay before the associated genes become transcriptionally active (**Fig. 2l**). The accessible state of these loci in the earliest progenitors may represent a form of priming to enable rapid cell-cycle entry once the decision to differentiate has been made and represents the type of discovery that can be enabled through integrative analysis across modalities.

### Robustness and benchmarking analysis

As our strategy relies on the ability for the dictionary to represent and reconstruct individual datasets, we explored how the size and composition of the multi-omic dataset affected the accuracy of integration. We sequentially downsampled the multi-omic dataset, repeated bridge integration, and compared the results with our original findings. Downsampling the bridge generally returned results that were concordant with the full analysis, but as expected, could affect annotation accuracy for rare cell types which are most sensitive to downsampling (**Fig. 3a**). We found that if a bridge dataset contained at least 50 cells (‘atoms’) representing a given cell type, this was sufficient for robust integration. We note that this threshold is not a strict requirement; we found that integration can be successful for rare cell types such as ASDC even when fewer than ten cells are present in the bridge, but we also observed failure modes in this regime. We note that generating bridge datasets consisting of more than 50 cells per subpopulation is quite feasible for many multi-omic technologies, and that our findings represent guidelines to assist in experimental design when performing multi-omic experiments.

**Figure 3.**
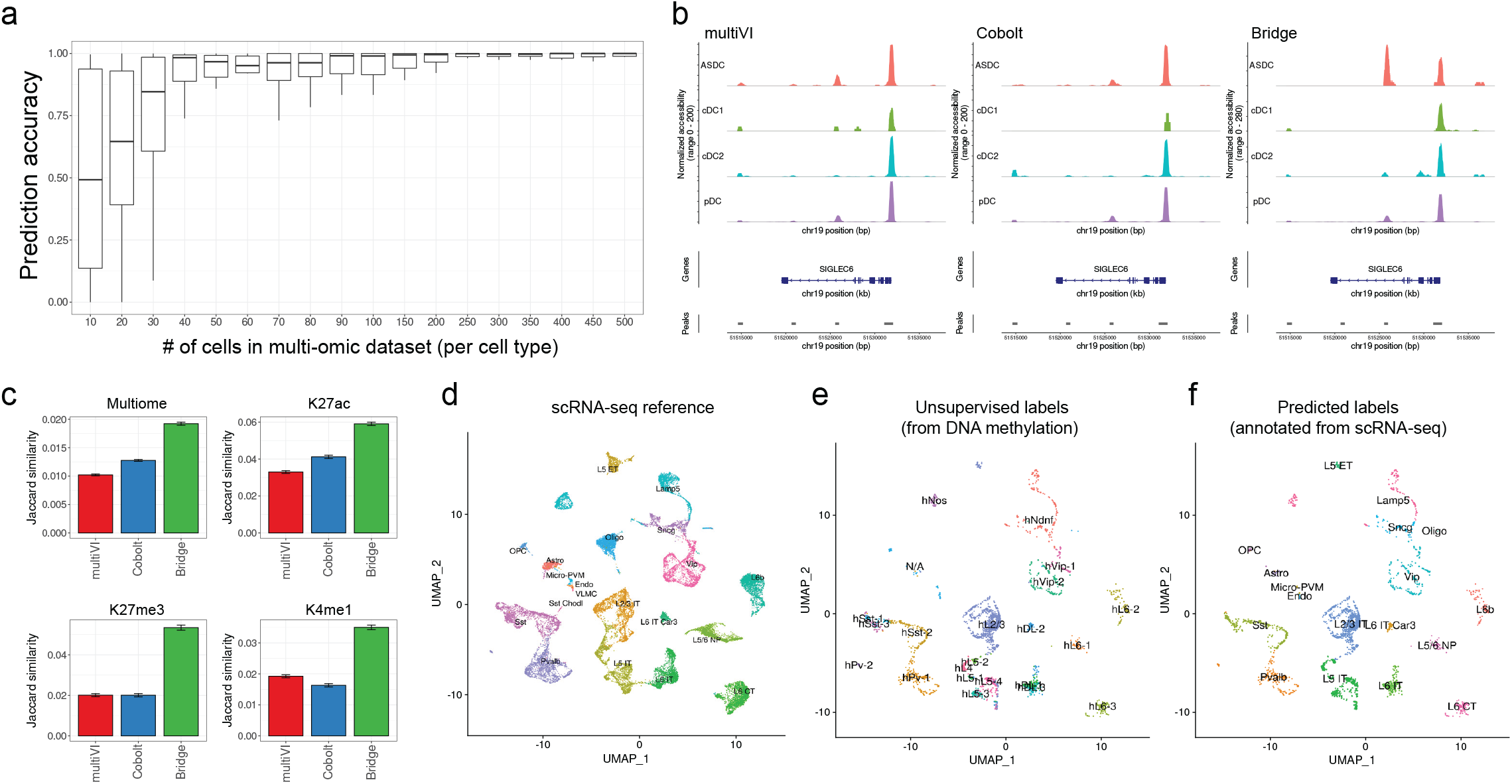
Robustness and benchmarking analysis for bridge integration. (**a**) Per cell-type prediction accuracy of bridge integration, based on the number of cells representing each cell type in the multi-omic dataset. Accuracy results were obtained by serially downsampling the multi-omic dataset, repeating bridge integration, and comparing resulting query annotations with those derived from the full dataset. Boxplots represent the observed range of values across 20 cell types. (**b**) Coverage plots for the SIGLEC6 locus, after performing crossmodality annotation with bridge integration, multiVI, and Cobolt. Only cells called as ASDC by bridge integration exhibit celltype-specific accessibility at this locus. Additional loci shown in Supplementary Fig. 2. (**c**) Ground truth benchmarking analysis. RNA and ATAC profiles from a 10x multiome dataset were unpaired and integrated. Barplots show the average Jaccard similarity between each scATAC-seq cell and its matched scRNA-seq cell. Results are split by individual cell types in Supplementary Fig. 2. Results are also shown for Paired-Tag datasets for three histone modification profiles. In each case, bridge integration achieves the highest Jaccard similarity. (**d**) scRNA-seq reference of the human motor cortex. (**e,f**) Mapping of single cell DNA methylation profiles of human cortical cells onto the reference using a snmC2T-seq multi-omic dataset as a bridge. Cells are colored by the methylation-derived annotations from the original study (**e**), or the scRNA-seq derived labels from bridge integration (**f**). Reference-derived labels at higher levels of granularity are shown in Supplementary Fig. 2.

We next compared the performance of bridge integration against two recently proposed methods for integrated analysis of multimodal and single-modality datasets. Both multiVI^48^ and Cobolt^49^ utilize variational autoencoders for integration, and while they do not explicitly treat multi-omic datasets as a bridge, they aim to integrate datasets across technologies and modalities into a shared space. When applied to the previously described datasets, both methods were broadly successful in integrating scRNA-seq and scATAC-seq data, but did not identify matches at the same level of resolution (for example, neither method successfully matched ASDC in scATAC-seq data to the ASDC in the Azimuth reference) (**Fig. 3b** and **Supplementary Fig. 2a-c**). When comparing computational efficiency, bridge integration (0.8 hours, not including 1.2 hours of preprocessing time), and Cobolt (3.3 hours) were the most efficient methods, while multiVI required more computational resources (15.7 hours).

To quantitatively benchmark performance, we split the 10x multi-omic bridge dataset into two groups. In one group, we treated the scRNA-seq and scATAC-seq data as if they were from separate experiments, representing a benchmark dataset for integration where ground-truth correspondences were known. The second group of cells was used as a multi-omic bridge dataset. After aligning cells across modalities, we calculated the Jaccard similarity metric between each scATAC-seq cell and its matched scRNA-seq counterpart. We found that our bridge integration strategy consistently maximized this similarity metric, demonstrating that our procedure most effectively matched cells in the same biological state across modalities (**Fig. 3c** and **Supplementary Fig. 2d**). Consistent with our previous results, we found that the strongest improvements were observed when mapping rare cell types including plasma cells and dendritic cells (**Supplementary Fig. 2d**). As our procedure is compatible with multiple integration techniques, we compared the performance of bridge integration when using either mnnCorrect^38^ or Seurat v3^19^ for the final alignment step, and observed very similar results (**Supplementary Fig. 2d**).

As a second quantitative benchmark with ground-truth data, we pursued a similar strategy using a recently published Paired-Tag dataset^26^, where individual histone modification binding profiles via scCUT&Tag were simultaneously measured with RNA transcriptomes. Since each Paired-Tag experiment was performed with biological replicates, we used one replicate as a multi-omic bridge dataset and split the other replicate into separate modalities for benchmarking. We performed cross-modality integration between scRNA-seq and scCUT&Tag for active histone marks (H3K27ac), repressive histone marks (H3K27me3), and enhancer histone marks (H3K4me1). In each case, bridge integration successfully integrated cells across modalities, and returned the highest Jaccard similarity between matched scRNA-seq and scCUT&Tag profiles (**Fig. 3c** and **Supplementary Fig. 2e-g**).

To further demonstrate the flexibility of our approach, we used bridge integration to map and annotate a snmC-seq dataset, which measures DNA methylation profiles in single cells from the human cortex^50^. As a reference, we utilized a dataset from the Allen Brain Atlas which defines a taxonomy of cell-types in the human cortex, and is accompanied by an expertly curated and multilevel cell ontology^51^. Using a snmC2T-seq dataset which simultaneously measures methylation and gene expression as a bridge^28^, we were able to annotate the snmC-seq profiles with high confidence (**Supplementary Fig. 2h**). Even when our reference-derived annotations did not augment the resolution to unsupervised clustering of snmC-seq data, they did add substantial interpretability (**Fig. 3d-f**). For example, unsupervised clustering identified multiple populations of L6 neurons (labeled as L6-1, L6-2, and L6-3), but RNA-assisted annotation clearly labeled these clusters as either ‘Near Projecting’ (NP) or deep neocortical laminar 6b (L6b) excitatory neurons (**Fig. 3f**).

Taken together, these results demonstrate the accuracy, robustness, and flexibility of our bridge integration procedure. We demonstrate applications on multiple modalities and data types, as well as best-in-class performance via quantitative and ground-truth benchmark comparisons. We demonstrate how cross-modality mapping can help interpret and improve the resolution of cell type annotation, including extremely rare cell types whose identification is facilitated by curated annotation in a reference dataset. Moreover, projecting datasets into a harmonized space also enables exploration of cross-modality relationships.

### Utilizing dictionary learning for scalable integration

The recent increase in publicly available single-cell datasets poses a significant challenge for integrative analysis. For example, multiple tissues have now been profiled across dozens of studies, representing hundreds of individuals and millions of cells. We refer to the challenge of harmonizing a broad swath (or the entirety) of publicly available single-cell datasets from a single organ as ‘community-wide’ integration. While a rich diversity of analytical methods can harmonize datasets of hundreds of thousands of cells, performing unsupervised ‘community-wide’ integration remains challenging, even when analyzing a single modality.

We were inspired by previous work on ‘geometric sketching’ which first selects a representative subset of cells (a ‘sketch’) across all datasets, integrates them, and then propagates the integrated result back to the full dataset^52,53^. This pioneering approach substantially improves the scalability of integration as the heaviest computational steps are focused on subsets of the data. However, this approach is dependent on the results of principal components analysis that must first be performed on the full dataset. As datasets continue to grow in scale, more sophisticated computational infrastructure is required to load full collections of data into memory, and even performing dimensional reduction can become a limiting step. We aimed to devise a strategy that could integrate large compendiums of datasets, without ever needing to simultaneously analyze or perform intensive computation on the full set of cells.

We reasoned that dictionary learning could also enable efficient and large-scale integrative analysis. We first select a representative sketch of cells (i.e. 5,000 cells) from each dataset, and treat these cells as atoms in a dictionary (**Fig. 4a,** Supplementary Methods). We next learn a dictionary representation, representing a weighted linear combination of atoms that can reconstruct the full dataset. These steps can occur for each dataset independently, allowing for efficient processing. We then perform integration on the atoms from each dataset. This is the only step that simultaneously analyzes cells from multiple datasets, but since only the atoms are considered, this does not impose scalability challenges. Finally, we apply our previously learned dictionary representations to the harmonized atoms from each dataset individually, and reconstruct harmonized profiles for the full dataset. We refer to this procedure as ‘atomic sketch integration’. We highlight that for this application, the ‘atoms’ used to reconstruct a dataset represent a subset of cells from the dataset itself. Contrastingly, in bridge integration, the atoms refer to cells from a different (multi-omic) dataset.

**Figure 4.**
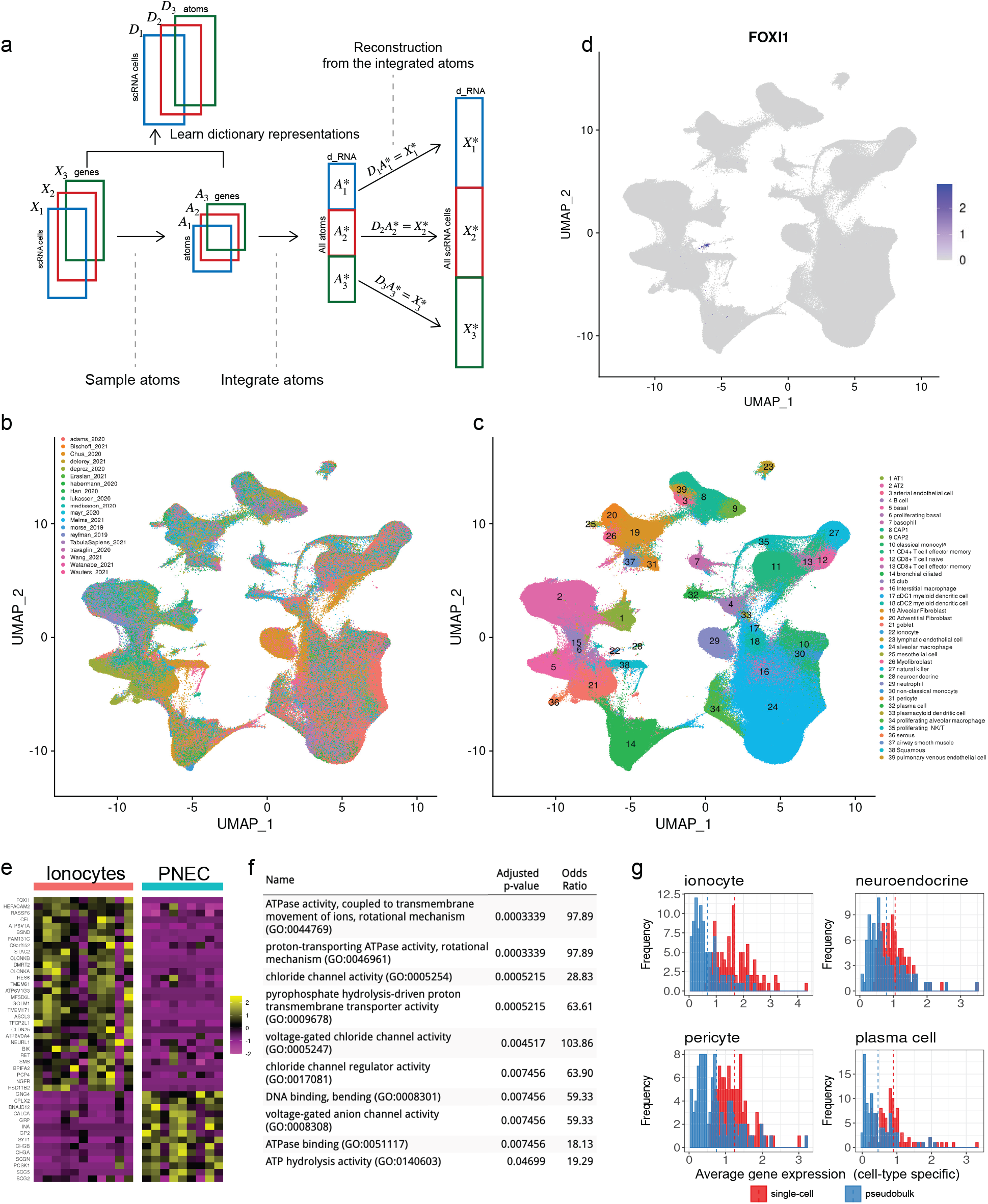
Utilizing dictionary learning for massively scalable integration. (**a**) Schematic of atomic sketch integration procedure. After selecting a representative set of cells from each dataset, these cells are integrated and used to reconstruct harmonized profiles for all cells. Matrix notation is consistent with the full mathematical description in Supplementary Methods. (**b, c**) UMAP visualization of 1,525,710 scRNA-seq profiles spanning 19 studies from the lung and upper airways, which were harmonized using atomic sketch integration in 55 minutes. Cells are colored by their study of origin (**b**) or annotated cell type after integration (**c**). (**d**) Expression of *FOXI1,* a transcriptional marker of pulmonary ionocytes, in the integrated dataset. (**e**) Heatmap showing the top transcriptional markers of pulmonary ionocytes that are consistent across multiple studies. Pulmonary neuroendocrine cells (PNEC), the most transcriptionally similar cell type, are shown for contrast. Each column represents a pseudobulk average of all cells from a single cell type and single study. Top transcriptional markers for all cell types are shown in Supplementary Fig. 4. (**f**) GO ontology enrichment terms for ionocyte markers. (**g**) Expression distributions of top transcriptional markers recovered from single-cell differential expression analysis (red), or pseudobulk analysis (blue).

The success of atomic sketch integration rests on identifying a representative subset of cells for each dataset. Sketching techniques for single-cell analysis aim to find subsamples that preserve the overall geometry of these datasets^52–54^. These methods do not require a pre-clustering of the data, but aim to ensure that the sketched dataset represents both rare and abundant cell states, even after downsampling. Here, we perform sketching using a leverage-score sampling based strategy that has been proposed for large-scale information retrieval problems^55^ and can be rapidly and efficiently computed on sparse datasets (Supplementary Methods). We emphasize that atomic sketch integration represents a general strategy for improving scalability that can be broadly coupled with existing methods. For example, a wide variety of integration techniques - including Harmony^37^, Scanorama^39^, mnnCorrect^38^, scVI^40^, and Seurat^19^, can be used to integrate the atom elements in each dictionary, with our procedure then enabling these results to be extended to full datasets.

### Community-scale integration for human lung scRNA-seq

To demonstrate the potential of atomic sketch integration to perform ‘community-wide’ analysis, we first considered scRNA-seq datasets of the human lung. During the COVID-19 pandemic, there has been widespread scRNA-seq data collection from respiratory tissues, particularly by the Human Cell Atlas Lung Biological Network^56^. Leveraging a recently published ‘database’ of scRNA-seq studies^57^, as well as collection of openly released lung and upper airway datasets from the Human Cell Atlas, we assembled a group of 19 datasets spanning 1,525,710 total cells. We created an atomic dictionary consisting of 5,000 cells from each dataset (95,000 total atoms), integrated these cells, and then reconstructed the full datasets. Our atomic sketch integration procedure performed all these steps (including preprocessing) in 55 minutes, using a single computational core.

Our results exhibit the advantages of community-scale integration compared to individual analysis. First, by matching biological states across datasets and technologies, the integrated reference can help to standardize cell ontologies and naming schemes (**Fig. 4b,c**). When observing previously assigned annotations derived from each study, we found that matched cell populations were often assigned slightly different names (**Supplementary Fig. 3a**). Unsupervised integration at this scale is a valuable tool for identifying these conflicts and can assist in the development of authoritative and standardized cell ontologies.

As a second benefit, we found that community-scale integration enabled consistent identification of ultra-rare populations, and in particular, a population of Foxi1-expressing ‘pulmonary ionocytes’ that were recently discovered in both human and mouse lungs^58^ (**Fig. 4d**). While these cells were only independently annotated in 6 out of 19 studies, our integrated analysis discovered at least one pulmonary ionocyte in 17 out of 19. The identified ionocytes were extremely rare (0.047%), but exhibited clear expression of canonical markers (**Fig. 4b**), highlighting the potential value for pooling multiple datasets to characterize these cells. We note that selection of dictionary atoms by sketching, or leverage-score sampling is essential for optimal performance (**Supplementary Fig. 3b,c)**; repeating the analysis using a set of atoms determined by random downsampling successfully integrated abundant cell types, but failed to integrate ionocytes as they were not sufficiently represented in the dictionary.

Finally, we found that community-scale integration can substantially improve the identification of differentially expressed (DE) cell-type markers. The use of 19 study replicates specifically enables us to identify genes that show consistent patterns across laboratories and technologies, representing robust and reproducible markers. We grouped cells by both sample replicate and cell type identity, and performed differential expression on the resulting pseudobulk profiles (**Fig. 4e** and **Supplementary Fig. 4**). For example, we identified 116 positive markers for pulmonary ionocytes, representing one of the deepest transcriptional characterizations of this cell type. These markers included both canonical markers such as the transcription factor *FOXI1,* but also revealed clear ontology enrichments for ATPases (e.g. *ATP6V1G3, ATP6V0A4)* and chloride channels (e.g. *CLCNKA, CLCNKB, CFTR),* supporting the role of these cells in regulating chemical concentrations in the lung (**Fig. 4f**). One advantage of working with pseudobulk values is increased quantification accuracy for lowly expressed genes. Indeed, we repeatedly found that top DE markers found using this strategy tended to capture more genes at a lower range of average expression values (**Fig. 4g**).

### Community-scale integration of scRNA-seq and CyTOF

As a final demonstration, we considered a similar problem of community-wide integration for circulating human peripheral blood cells, which is one of the most widely profiled systems with diverse single-cell technologies. Exploring publicly available studies of either COVID-19 samples or healthy controls, we accumulated a collection of 14 studies with scRNA-seq measurements, representing a total of 3.46M cells from 639 individuals. Data from 11 of the studies was obtained from a recently published collection of standardized single-cell sequencing datasets^59^. We performed unsupervised atomic sketch integration, yielding a harmonized collection in which we annotated 30 cell states (**Fig. 5a**). As a subset of our samples were not depleted for granulocytes, our collection includes a distinct population of neutrophils that were absent in our previous Azimuth reference of human PBMC. Moreover, we identified specific populations of activated granulocytes and B cells that were specific to COVID-19 samples (**Supplementary Fig. 5a**). Consistent with previous reports, monocytes in COVID-19 samples sharply upregulated interferon response genes^60,61^, but were correctly harmonized with healthy monocytes (**Fig. 5b** and **Supplementary Fig. 5b**). By matching shared cell types across disease states (while still allowing for the possibility of disease-specific subpopulations), this collection represents a valuable resource for identifying cell-type specific transcriptional changes that reproduce across multiple studies. We characterized cell-type specific responses for eight additional cell types, each of which exhibited a conserved interferon-driven response alongside the activation of cell typespecific response genes (**Supplementary Fig. 6**).

**Figure 5.**
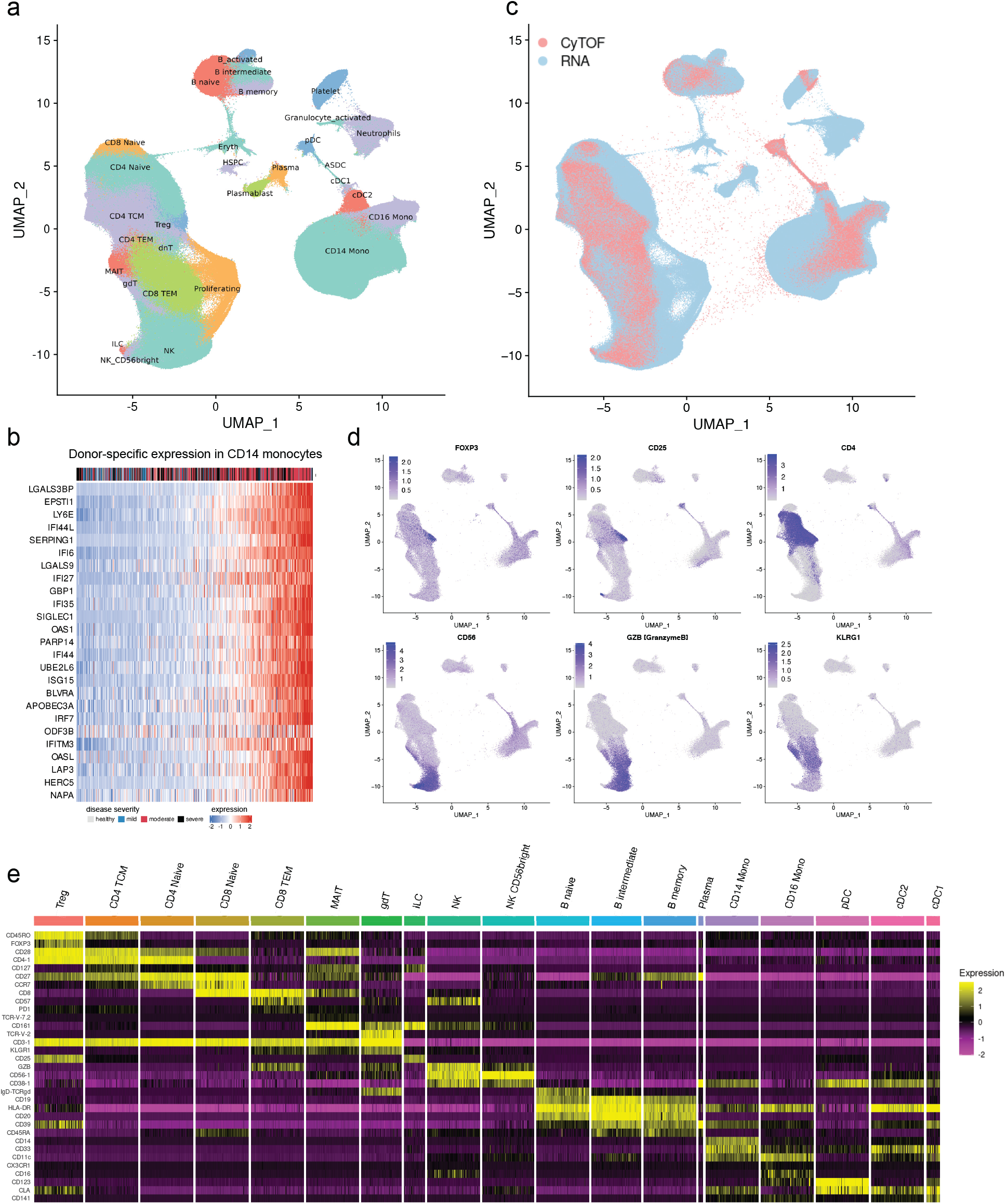
‘Community-scale’ integration of sequencing and cytometry immune datasets. (**a**) UMAP visualization of 3,461,171 human PBMC scRNA-seq profiles spanning 14 studies and 639 individuals after performing atomic sketch integration. (**b**) Expression of a COVID-19 response module in CD14 monocytes. Each column represents a pseudobulk average of CD14 monocytes from one of 506 individuals. Expression of the module is correlated with disease severity within the individual, which is indicated by the color scale above the heatmap. Responses for additional cell states are shown in Supplementary Fig. 5b. (**c**) Mapping of 5,170,249 additional CyTOF profiles spanning 119 individuals, using a published CITE-seq dataset (Hao et al, 2021) as a multi-omic bridge. Each CyTOF profile is annotated with one of the scRNA-seq defined cell types. (**d**) Cross-modality integration enables the exploration of cell surface and intracellular protein markers on cell landscapes defined by scRNA-seq. As an example, intracellular FOXP3 levels are highly enriched in annotated Treg cells, validating the accuracy of our mapping. 200,000 cells are shown in each visualization to alleviate overplotting. (**e**) Heatmap showing the expression of 34 protein markers in the CyTOF dataset. Each column represents a pseudobulk average, after grouping cells by individual and reference-derived annotation.

While single-cell sequencing technologies are capable of measuring RNA transcripts and surface proteins in thousands of single cells, cytometry-based techniques can measure both extracellular and intracellular proteins in millions of cells. As our bridge integration procedure should enable the mapping of CyTOF profiles onto scRNA-seq datasets, we obtained a collection of CyTOF datasets spanning 119 individuals and 5,170,249 total cells^62^. We used our previously collected CITE-seq dataset of 161,764 PBMC from healthy donors as a multi-omic bridge^4^. The CyTOF and CITE-seq dataset both shared 30 cell surface protein features, while the CyTOF dataset also measured 17 unique proteins which included intracellular targets that cannot be measured via CITE-seq.

Bridge integration annotated each CyTOF dataset with cluster labels derived from our 3.46M cell scRNA-seq collection, and allowed us to infer intracellular protein levels for each of these clusters (**Fig. 5c**). Predicted regulatory CD4^+^ T cells expressed high levels of the transcription factor Foxp3^63^, and effector T cells exhibited enriched Klrg1 levels^64^ (**Fig. 5d**). We also found that among cytotoxic lymphocyte populations, MAIT cells were uniquely depleted for expression of the cytotoxic protease Granzyme B, consistent with previous reports^65^. Each of these patterns supports the accuracy of our cross-modality mapping. Finally, we successfully annotated a rare populations of innate lymphoid cells (0.024%), which were not independently identified in the CyTOF dataset, but correctly exhibited a CD25^+^CD127^+^CD161^+^CD56^-^ immunophenotype^4,66^ (**Fig. 5d,e**). Taken together, we conclude that dictionary learning enhances the scalability of integration, as well as the ability to integrate and compare diverse molecular modalities.

## Discussion

In order to map datasets measuring a diverse set of modalities to scRNA-seq reference datasets, we developed bridge integration, an approach for cross-modality alignment that leverages a multiomic dataset as a bridge. We characterize specific compositional requirements for the bridge dataset, perform quantitative benchmarking analyses with ground-truth datasets, and demonstrate the broad applicability of our method to a wide variety of technologies and modalities. Finally, we demonstrate how to use atomic sketch integration to extend the scalability of our approach to harmonize dozens of datasets spanning millions of cells.

We anticipate that our methods will be valuable to both individual labs but also larger consortia that have already invested in constructing and annotating comprehensive scRNA-seq references. For example, the Human Cell Atlas, Human Biomolecular Atlas Project, Tabula Sapiens^67^, and Human Cell Landscape^68^, all have released scRNA-seq references spanning hundreds of thousands of cells for multiple human tissues. Similar efforts are present in model organisms as well, including the Fly Cell Atlas^69^, and Plant Cell Atlas projects^70^. In each case, these efforts involve careful, collaborative, and expert-driven cell annotation alongside the curation of reference cell ontologies. While repeating this manual effort for each modality is infeasible, bridge integration enables the mapping of new modalities without having to modify the reference. As additional multi-omic datasets become available, we expect that tools such as Azimuth will begin to map additional modalities as well.

We note that the bridge integration is particularly well-suited for experimental designs where multiomic technologies can be applied to a subset, rather than all, experimental samples. This is a common occurrence, particularly because multi-omic technologies often are associated with increased cost, lower throughput, and reduced data quality for each individual measurement type. In particular, we note that combinatorial indexing approaches can be readily applied using commercial instrumentation to profile a single modality in hundreds of thousands of cells^71,72^, but the same is not true for multi-omic technologies. We propose that the collection of large single-modality datasets, harmonized via a smaller but representative multi-omic bridge, may represent an efficient and robust strategy to explore cross-modality relationships across millions of cells. Our identification of cell cycle ‘priming’ in hematopoietic stem cells represents an example of cross-modality insights that can be derived via bridge integration.

We note that future extensions of our work can further broaden the applicability of bridge integration or demonstrate its potential in new contexts. For example, performing bridge integration on spatially resolved unimodal datasets (e.g. CODEX^73^), could help to better characterize the spatial localization of scRNA-seq defined cell types in large tissue sections. New multi-omic technologies that couple high-resolution mass spectrometry imaging to single-cell or spatial transcriptomics could serve as a bridge to harmonize lipidomic and metabolic profiles^74,75^ with sequencing-based references. In addition, future computational improvements will further lower the requirements of the bridge dataset, enabling robust integration with an even smaller number of multi-omic cells.

We emphasize the ability for bridge and atomic sketch integration to identify and characterize rare cell populations, including AXL^+^ SIGLEC6^+^ dendritic cells and pulmonary ionocytes. Single cell transcriptome profiling played an essential role in the initial discovery of these cell types, but a deeper understanding of their biological role and function will benefit from multimodal characterization. The goal of moving beyond an initial taxonomic classification of cell types towards a complete multimodal reference will not be accomplished with a single experiment or technology. We envision that computational tools for cross-modality integration will play key contributions to the construction of this map.

## Author Contributions

TS, YH, and RS conceived the research. YH, TS, MK, SC, PH, AH, AS, GM, SM performed computational analysis, supervised by CG and RS. YH TS and RS wrote the manuscript, with input and assistance from all authors.

## Acknowledgements

The authors would like to thank all the members of the Satija Lab for thoughtful discussions related to this work. We thank Andrew Butler and Harsh Srivastava for assistance in identifying and locating scRNA-seq datasets from human lung and PBMC. We acknowledge the Gottardo and Newell labs for publicly releasing a standardized compendium of human PBMC scRNA-seq datasets. This work was supported by the Chan Zuckerberg Initiative (EOSS-0000000082, HCA-A-1704-01895 to R.S.), and the NIH (K99HG011489-01 to T.S; K99CA267677 to A.S; RM1HG011014-02, 1OT2OD026673-01, DP2HG009623-01, R01HD096770, R35NS097404 to R.S).

## Competing Interests

In the past three years, R.S. has worked as a consultant for Bristol-Myers Squibb, Regeneron, and Kallyope and served as an SAB member for ImmunAI, Resolve Biosciences, Nanostring, and the NYC Pandemic Response Lab. The other authors declare no competing interests.

## Data Availability

We used publicly available datasets in this work. Download locations for each dataset are listed in the Supplementary Methods, and also Supplementary Table 1. Azimuth references are available for download at http://azimuth.hubmapconsortium.org

## Code Availability

Bridge Integration and Atomic Sketch Integration are implemented as part of the Seurat R package. In this work, we also make use of the Signac, and Azimuth packages. All are freely available as open-source software:

https://github.com/satijalab/seurat

https://github.com/timoast/signac

https://github.com/satijalab/azimuth

## Supplementary Figure Legends

**Supplementary Figure 1:**
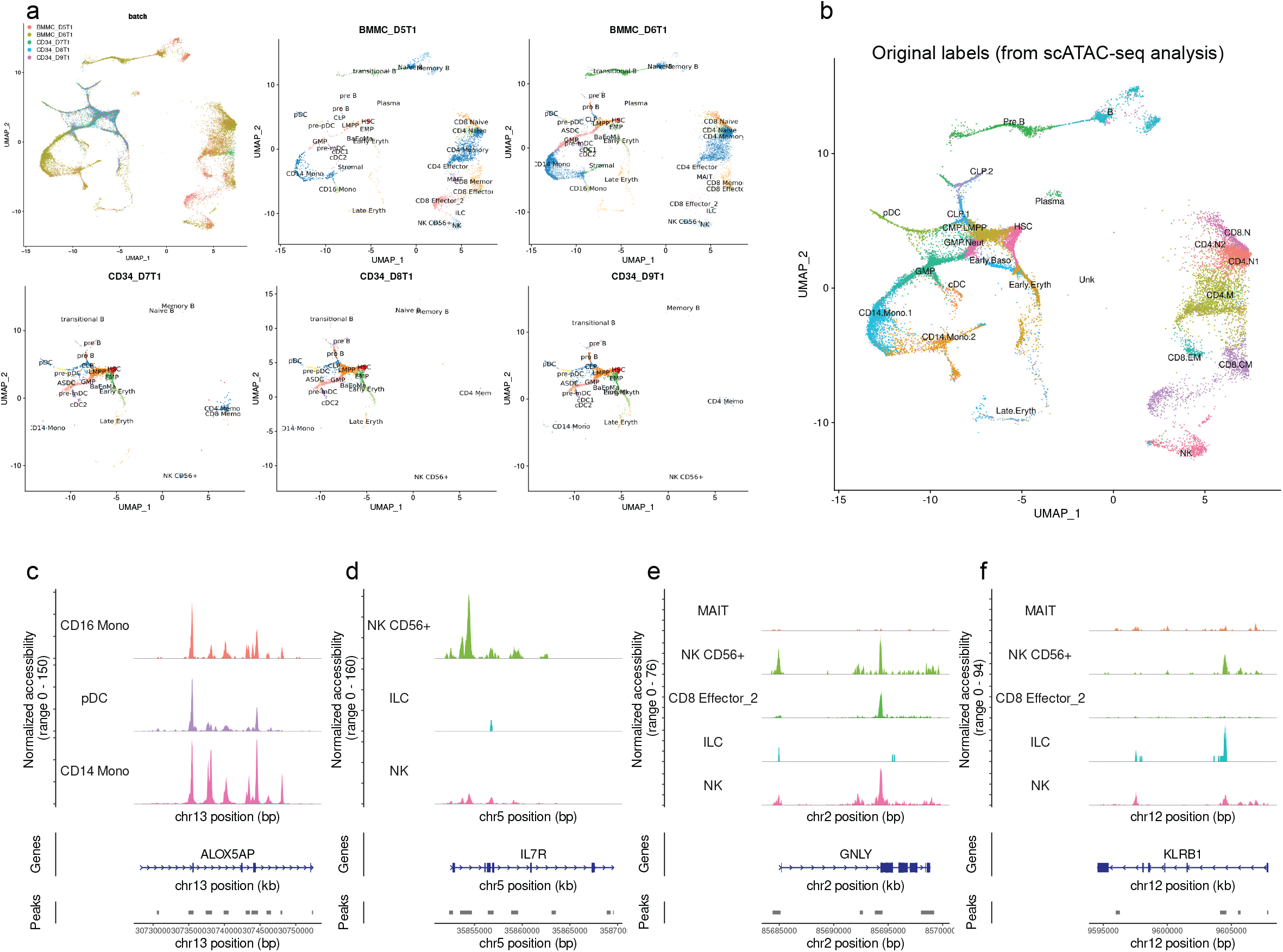
Mapping scATAC-seq data onto scRNA-seq references. Related to Figure 2. (a) UMAP visualization of an scATAC-seq query dataset from (Granja et al, 2019), representing 26,159 profiles across 5 samples. Same as Fig. 2c, but in the top-left panel, cells are colored by sample identity. Remaining panels show cells from the individual batches, including three batches (bottom row) that are enriched for CD34^+^ cells. Cells are colored by their reference-derived annotation after bridge integration. Enriched samples overwhelmingly map to progenitor populations in the reference map. (b) Same as (a), but cells are colored by their unsupervised annotations assigned by the authors of (Granja et al, 2019). The annotations are generally concordant, but reference-derived annotations provide additional granularity and help to resolve ambiguous cell names, including for CD16^+^ monocytes, CD56^+^ NK cells, Innate Lymphoid Cells (ILC), and Mucosal Associated Invariant T (MAIT) cells. (c-f) Coverage plots showing chromatin accessibility at selected loci, after grouping query cells by their predicted annotations. In each case, the predicted cell labels agree with the expected accessibility patterns.

**Supplementary Figure 2.**
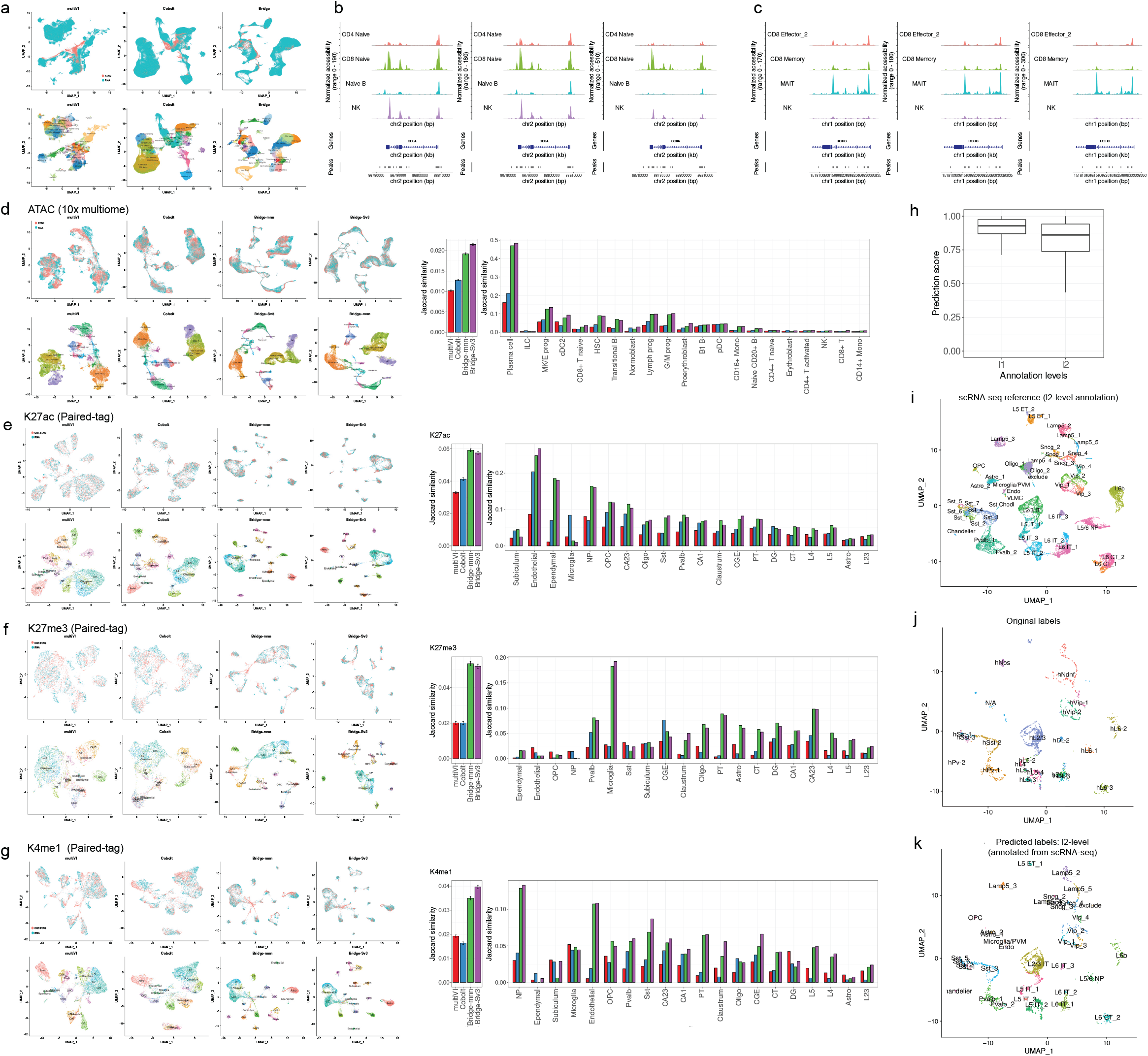
*Robustness and benchmarking analysis for bridge integration.* Related to Figure 3. (a) UMAP visualization of the mapped scATAC-seq dataset in Fig. 2, computed using the multiVI and Cobolt, and Bridge integration approaches. In Fig. 2, we visualize the results of bridge integration on the scRNA-seq reference-derived UMAP. Here, we compute a new UMAP visualization to enable comparison with alternative methods. (b) Since we do not have ground truth of cell labels for this dataset, we visualize chromatin accessibility patterns to assess annotation accuracy. At the CD8A locus, bridge integration shows the clearest celltype-specific accessibility patterns, suggesting that alternative methods blend CD8 and CD4 naïve cell populations together during integration. (c) At the RORC locus, all methods correctly infer MAIT-specific accessibility. (d) Ground-truth benchmarking, where integrated BMMC scRNA-seq and scATAC-seq profiles were originally measured in the same cells in a 10x multiome experiment. Left panel shows UMAP visualizations computed by all methods, with cells colored by either their measurement modality or expert-assigned cell annotation provided with the dataset. Right panel shows Jaccard similarities of matched cell profiles, either averaged across all cells, or split by cell annotation. We show results for Bridge integration using two alignment strategies for the final step, mnnCorrect (mnn), and Seurat v3 (Sv3). (e-g) Same as in (d), but ground-truth datasets originate from the Paired-Tag technology which measured individual histone modifications with cellular transcriptomes in the mouse brain. (h) Prediction scores for annotating single cell methylation profiles from an scRNA-seq reference via bridge integration. (i-k) Same as Fig. 3d-f, but with a higher level of granularity for the scRNA-seq cell annotations.

**Supplementary Figure 3.**
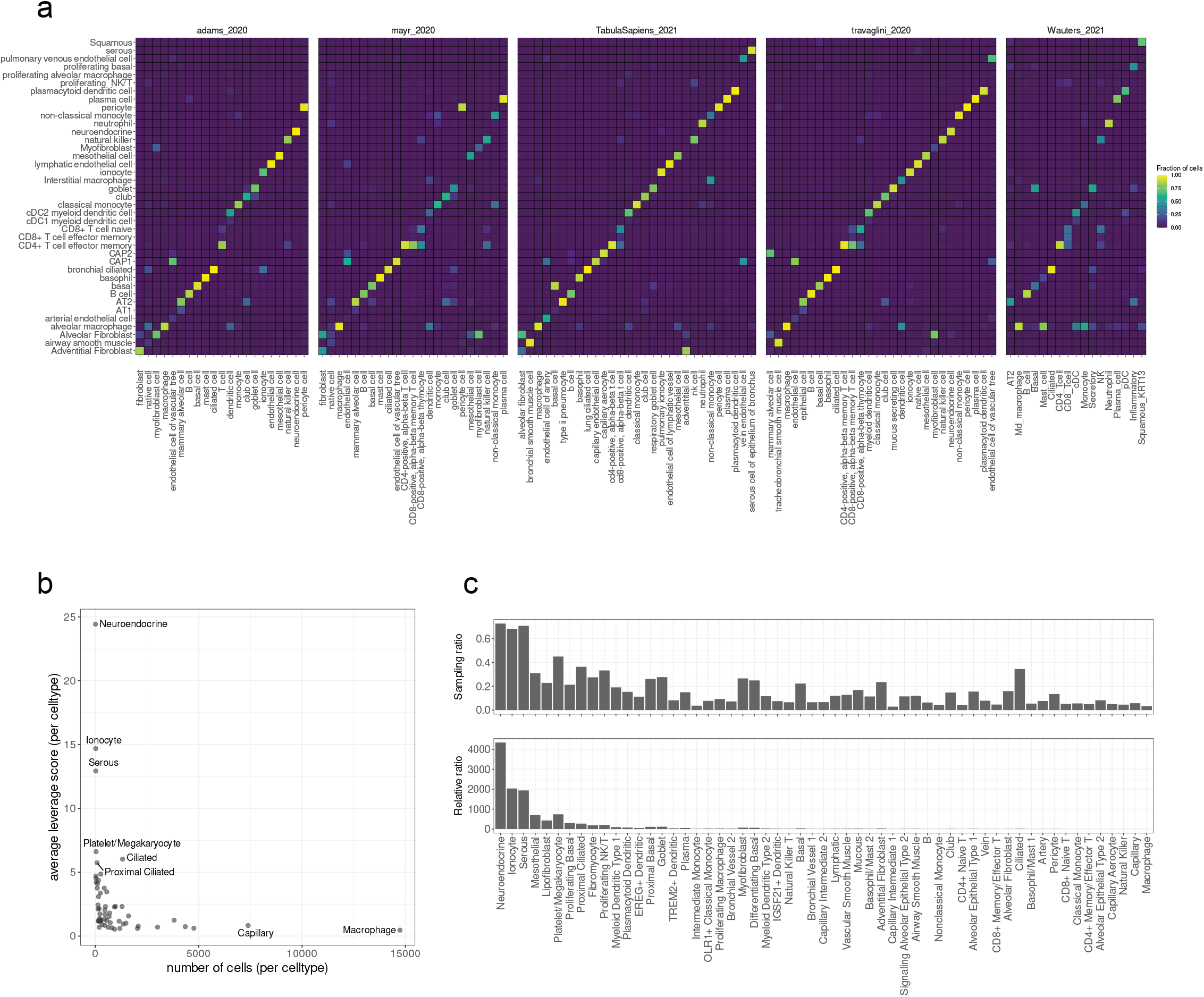
*Community-scale integration in the human lung*. Related to Figure 4. (a) Confusion matrices for five representative studies, showing the agreement between the originally assigned cell annotations (provided with each individual study), with the assigned annotation from our ‘community-wide’ integrative analysis of 1,525,710 scRNA-seq profiles. Large-scale integrative analyses can help to identify matches between disparate cell naming schemes and ontologies. (b) We calculated a ‘leverage score’ (Supplementary Methods) for each cell in each dataset, prior to performing any clustering analysis. Cells with high leverage scores should originate from rare populations, which is what we observe. Scatter plot shows relationship between the abundance of each cell population and the average leverage score all cells. (c) We used the leverage scores to sample a 5,000 cell ‘sketch’ from each dataset as atoms. Top barplot shows the probability of selecting cells from each population as an atom. For example, despite representing less than 0.05% of all cells in the dataset, we selected more than 60% of ionocytes as atoms. Bottom plot shows the relative enrichment of each cell population amongst the atoms, compared to the full dataset. Plots in (b-c) are shown for one representative dataset (tragavlini_2020).

**Supplementary Figure 4.**
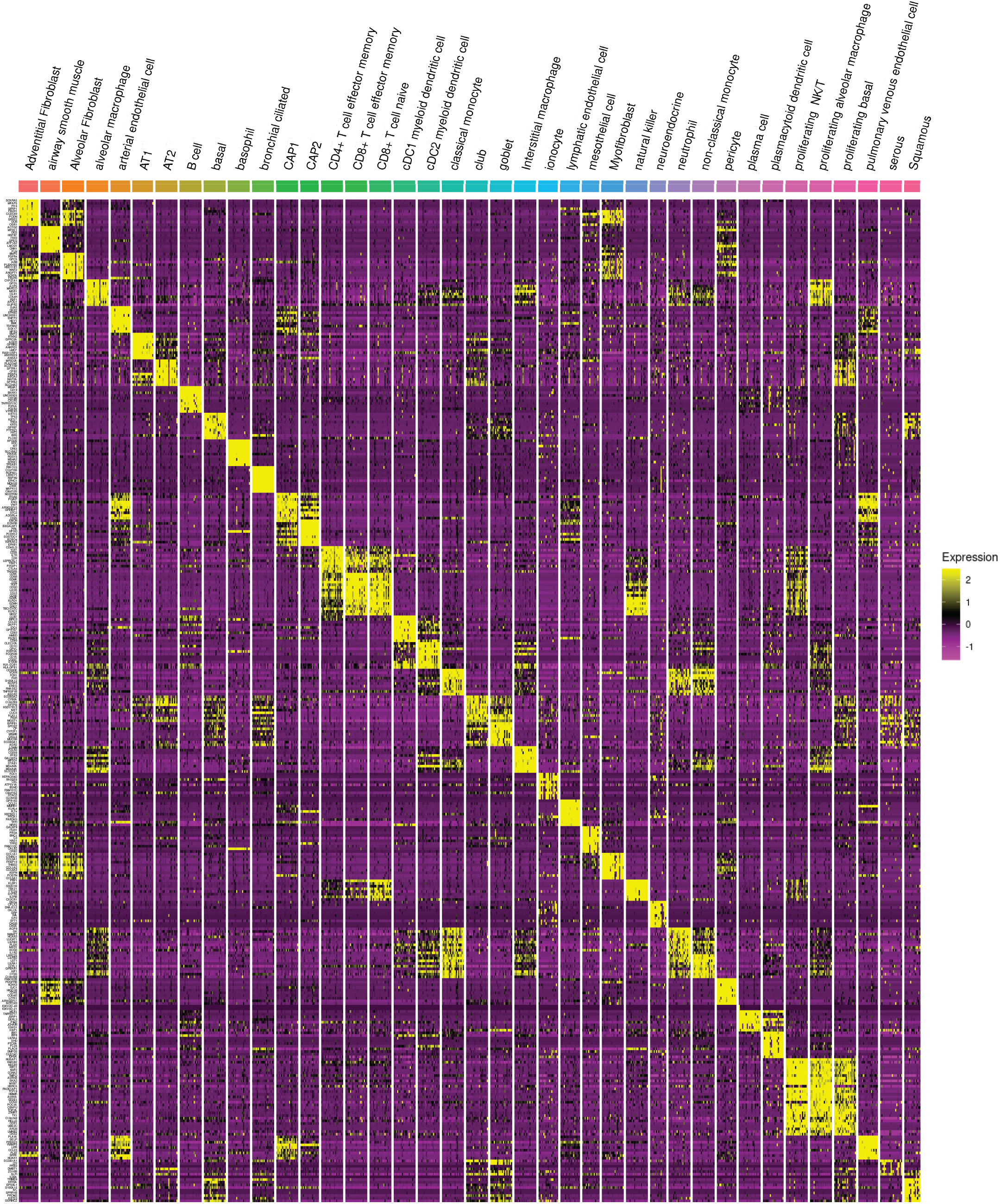
Heatmap of reproducible gene expression markers in human lung cell states. Related to Figure 4. Heatmap showing the top transcriptional markers of 39 cell states that are consistent across multiple studies. Each column represents a pseudobulk average of all cells from a single cell type and single study. The top ten markers are shown for each cell state.

**Supplementary Figure 5.**
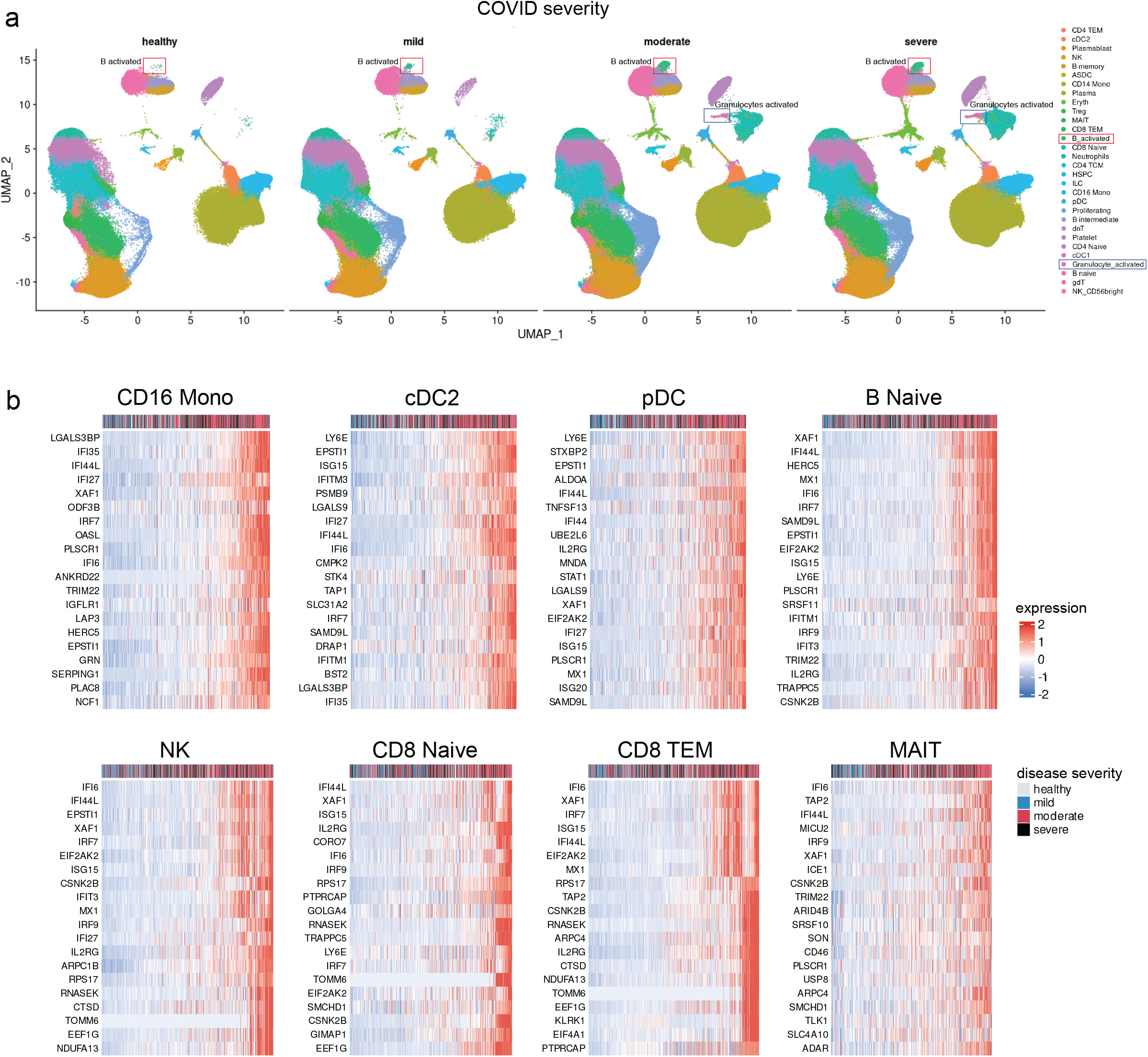
*Community-scale integration of sequencing and cytometry datasets*. Related to Figure 5. (a) UMAP visualization of integrated scRNA-seq profiles from human PBMC. Same as in Fig. 5a, but split into cells from healthy controls, or samples associated with mild, moderate, and severe COVID. Activated B cells are at sharply reduced frequency in the healthy control samples (which were also depleted for granulocytes). Activated granulocytes are observed primarily in moderate and severe COVID samples. (b) Same as Fig. 5b, but showing cell type-specific responses that correlate with disease severity for eight additional cell states. Each column represents a pseudobulk average from one of 506 individuals. Expression of the module is correlated with disease severity within the individual, which is indicated by the color scale above the heatmap.

**Supplementary Figure 6.**
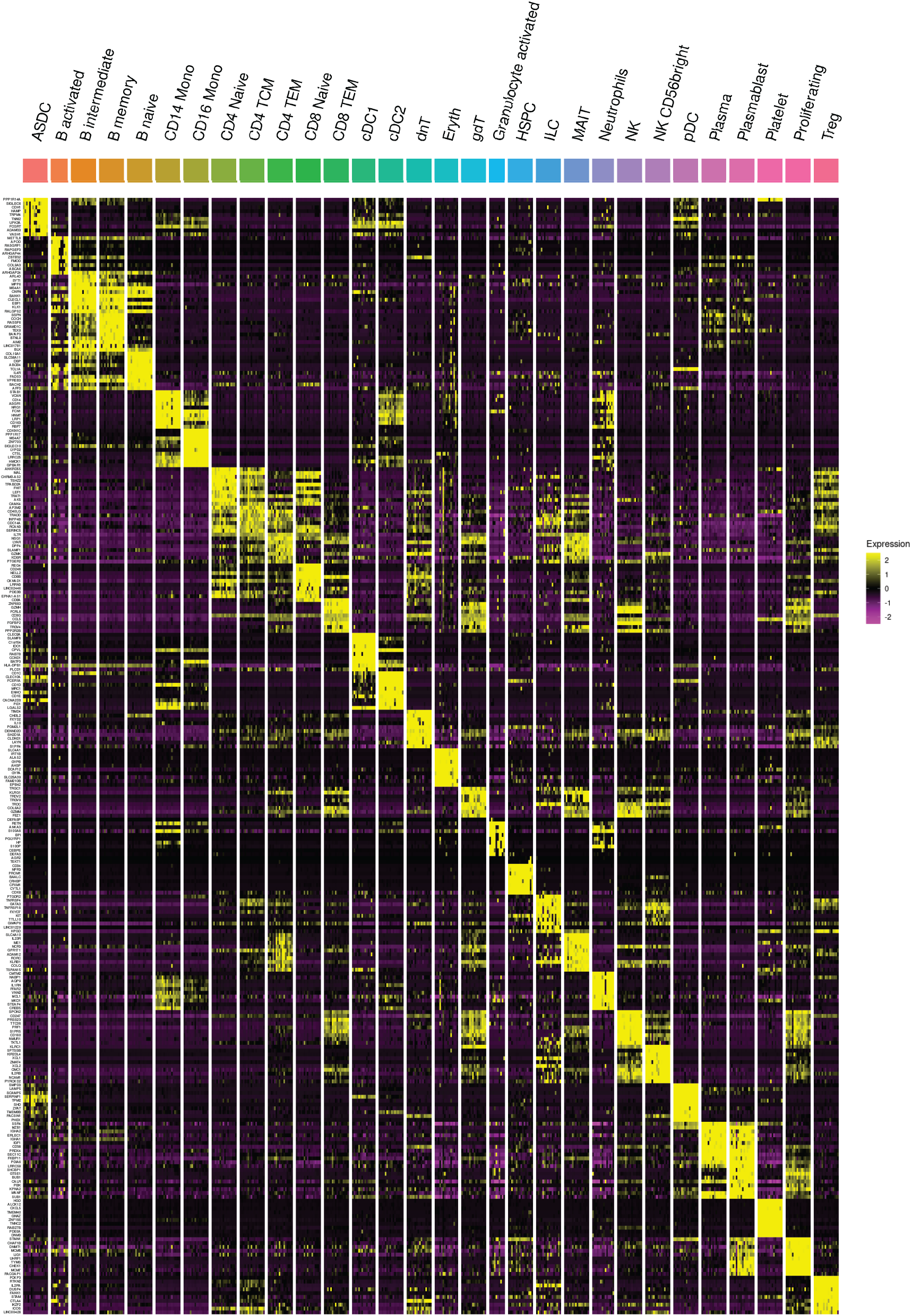
Heatmap of reproducible gene expression markers in human PBMC cell states. Related to Figure 5. Heatmap showing the top transcriptional markers of 30 cell states that are consistent across multiple studies. Each column represents a pseudobulk average of all cells from a single cell type and single study. The top ten markers are shown for each cell state.

