## Supplementary Methods for "Dictionary learning for integrative, multimodal, and scalable single-cell analysis"

**Bridge integration procedure**

Our bridge integration procedure is designed to perform integration of single-cell datasets profiling different modalities by leveraging a separate multi-omic dataset as a molecular bridge. The individual multi-omic profiles each represent individual atoms, which together comprise a multi-omic dictionary (i.e., each cell in the bridge dataset represents an atom, and the entire bridge dataset represents a dictionary). This dictionary is used to transform both unimodal datasets into a shared space defined by the same set of features, facilitating cross-modality integration. Our approach consists of four broad steps, described in detail below: (1) within-modality harmonization of unimodal and bridge datasets (2) Constructing a dictionary representation for each unimodal dataset (3) Dimensional reduction via Laplacian eigen decomposition. (4) Align dictionary representations across datasets. We illustrate each step of the method in Fig. 1b, using the same mathematical notation as we introduce below.

All methods are implemented in our open-source R package Seurat (www.satijalab.org/seurat, www.github.com/satijalab/seurat)

<u>1. Within-modality harmonization of unimodal and bridge datasets</u>

The first step in our procedure is to harmonize the unimodal and bridge datasets, based on shared modalities. For example, when performing bridge integration to map a scATAC-seq dataset onto an scRNA-seq reference (via a 10x multiome bridge), we first harmonize the gene expression measurements from the scRNA-seq and multiome experiments, and the chromatin accessibility measurements from the scATAC-seq and multiome experiments. Specifically, we define:

$X \in \mathbb{R}^{n_{scRNA-seq} \times d_{genes}}$: scRNA-seq expression counts matrix

$Y \in \mathbb{R}^{n_{scATAC-seq} \times d_{peaks}}$: scATAC-seq accessibility counts matrix

$M = [M_X M_Y]$: multi-omic expression + accessibility counts matrix, where

$M_X \in \mathbb{R}^{n_{multi-omic} \times d_{genes}}$: scRNA-seq subset of multi-omic matrix

$M_Y \in \mathbb{R}^{n_{multi-omic} \times d_{peaks}}$: scATAC-seq subset of multi-omic matrix

Our goal is to harmonize $X$ and $M_X$, and $Y$ and $M_Y$. This can be performed with a wide variety of existing tools for the harmonization of single-cell datasets. For example, Seurat, Harmony, LIGER, scVI, Scanorama, fastMNN, scVI, and scArches, all learn a shared low-dimensional space that jointly represents the datasets and aligns cells in a matched biological state. Our goal is therefore to learn:

$X^* \in \mathbb{R}^{n_{scRNA-seq} \times d_{RNA}}$: harmonized space for scRNA-seq data

$Y^* \in \mathbb{R}^{n_{scATAC-seq} \times d_{ATAC}}$: harmonized space for scATAC-seq data

$M^* = [M_X^* M_Y^*]$, where

$M_X^* \in \mathbb{R}^{n_{multi-omic} \times d_{RNA}}$: harmonized space for scRNA-seq subset of multi-omic dataset

$M_Y^* \in \mathbb{R}^{n_{multi-omic} \times d_{ATAC}}$: harmonized space for scATAC-seq subset of multi-omic dataset

In this work, we treat the scRNA-seq dataset $X$ as a reference and map the multi-omic gene expression profiles ($M_X$) onto this reference using the FindTransferAnchors and MapQuery functions in Seurat, in order to obtain $X^*$ and $M_X^*$. An example workflow is provided at (https://satijalab.org/seurat/articles/integration_mapping.html; 'Mapping and Annotating Query Datasets').

The same functionality has been implemented in the Signac package for the mapping and harmonization of scATAC-seq datasets (https://satijalab.org/signac/articles/integrate_atac.html). However, we emphasize that our approach is compatible with a wide variety of pre-existing approaches for within-modality harmonization, including all the methods listed above.

We also note that when finding anchors between the bridge and query datasets, we can leverage the multimodal nature of the bridge dataset to perform 'supervised' dimensional reduction, which utilizes both modalities when calculating a low-dimensional representation during harmonization. For example, we have previously described the use of 'supervised PCA' to learn optimized transformations from CITE-seq data[1,2]. When working with bridge datasets that measure ATAC-seq or CUT&Tag chromatin features (e.g. Paired-Tag, 10x multiome), we utilize an analogous procedure for supervising the latent semantic indexing (LSI) reduction.

2. Constructing a dictionary representation for each unimodal dataset

The goal of dictionary learning is to reconstruct individual data points as a weighted linear combination of atoms in a dictionary. We treat $M^*$ as a dictionary, with each row of this matrix representing an atom. We aim to learn reconstructions of $X^*$ and $Y^*$ based on the atoms of $M^*$, while minimizing the error between the original and reconstructed values. Specifically, we aim to identify the matrices $D_X$ and $D_Y$ where:

$D_X \in \mathbb{R}^{n_{scRNA-seq} \times n_{multi-omic}}$: dictionary representation of scRNA-seq dataset

$D_Y \in \mathbb{R}^{n_{scATAC-seq} \times n_{multi-omic}}$: dictionary representation of scATAC-seq dataset

Such that:

$$\underset{D_X}{argmin}( \|D_X(M_X^*) - X^*\|_F^2 + \|D_X\|_F^2)$$

and

$$\underset{D_Y}{argmin}( \|D_Y(M_Y^*) - Y^*\|_F^2 + \|D_Y\|_F^2)$$

As described in [3,4], this optimization problem is analogous to matrix regression, and has a closed-form solution for calculating $D_X$ and $D_Y$

$$D_X = X^*(M_X^*)^{\dagger}$$

$$D_Y = Y^*(M_Y^*)^{\dagger}$$

Where † represents the pseudoinverse of the matrix.

We note that $D_X$ and $D_Y$ represent transformations of the original scRNA-seq and scATAC-seq datasets. While the two experiments originally measured different sets of features, after the transformation, they now are represented by the same set of features, namely, the atoms of the multi-omic experiment.

3. Dimensional reduction via Laplacian Eigendecomposition

After the datasets have been transformed in the previous step, it is possible to integrate them directly. The dimensionality of the datasets is based on the number of cells in the multi-omic dataset. Unlike the original measurements, the dictionary representations are not sparse. As multi-omic datasets often consist of thousands of cells, working with high-dimensional and non-sparse is computationally inefficient. We therefore aimed to reduce the dimensionality of the dictionary representation. Motivated by a similar problem addressed by Laplacian Eigenmaps[5], a non-linear dimensionality reduction technique, we perform dimensionality reduction by computing an eigen decomposition of the graph Laplacian matrix.

We first compute a graph representation of the multi-omic dataset $M^*$. We utilize a 'shared nearest-neighbor' graph representation, as proposed for clustering single-cell datasets by[6]. We note that the matrix representation of this graph is symmetric, which is a requirement for downstream eigen decomposition. Our approach is compatible with any user-defined distance metric when constructing this graph, though we recommend utilizing either the Euclidean distance based on harmonized gene expression measurements (i.e $M_X^*$ ), or alternately, a weighted combination of modalities using the 'weighted-nearest neighbor' distance metric that we have previously introduced[2]. We define:

$G \in \mathbb{R}^{n_{multi-omic} \times n_{multi-omic}}$: Symmetric graph representation of multi-omic dataset

$L = I - D^{-\frac{1}{2}} G D^{-\frac{1}{2}}$ : Graph Laplacian matrix

We next perform an eigen decomposition of the Graph Laplacian matrix

$L = U \Lambda U^T$

$0 = \lambda_1 \leqslant \lambda_2 \leqslant \ldots \leqslant \lambda_n$

$U_L$: The leftmost $n_{laplacian}$ eigenvectors of $U$ , where *n* specifies the reduced dimensionality of the dataset. We select $n_{laplacian} = 50$ for all examples in this work.

We now multiply the learned dictionary representations for the scRNA-seq and scATAC-seq datasets by this truncated set of eigenvectors. Doing so transforms these representations into the same lower-dimensional space ($n_{laplacian}$). We define:

$L_X \in \mathbb{R}^{n_{scRNA-seq} \times n_{laplacian}}$: reduced dictionary representation for scRNA-seq data

$L_Y \in \mathbb{R}^{n_{scATAC-seq} \times n_{laplacian}}$ : reduced dictionary representation for scATAC-seq data

$L_M \in \mathbb{R}^{n_{multi-omic} \times n_{laplacian}}$: reduced dictionary representation for multi-omic dataset

And calculate these matrices:

$L_X = D_X U_L = X^* ( ( M_X^* )^{\dagger} U_L )$

$L_Y = D_Y U_L = Y^* ( ( M_Y^* )^{\dagger} U_L )$

$L_M = U_L$

4. Align dictionary representations across datasets.

Both the scRNA-seq and scATAC-seq dataset have now been transformed into a low-dimensional space defined by the same set of features. They can now be directly harmonized using existing methods. As in step (1), multiple published methods can accomplish this goal. In this work, we use our internal implementation of the mnnCorrect integration technique to perform this harmonization[7]. We choose mnnCorrect as we find that after performing the steps above, any remaining sample-specific differences are minor, and are typically far less than the differences we observe when aligning scRNA-seq datasets across different technologies. To demonstrate the compatibility of our approach with alternative methods, we also repeat our quantitative benchmarking experiments using our previously developed integration workflow in Seurat v3[8], and observe very similar results. (Supplementary Fig. 2).

Specifically, the final output of our procedure represents

$L_X^* \in \mathbb{R}^{n_{scRNA-seq} \times n_{laplacian}}$: harmonized reduced dictionary representation for scRNA-seq data

$L_Y^* \in \mathbb{R}^{n_{scATAC-seq} \times n_{laplacian}}$ : harmonized reduced dictionary representation for scATAC-seq data

$L_M^* \in \mathbb{R}^{n_{multi-omic} \times n_{laplacian}}$: harmonized reduced dictionary representation for multi-omic dataset

These representations can be used as input for common downstream analytical tasks including tSNE or UMAP visualization, graph-based clustering, and the identification of developmental trajectories.

**Atomic sketch integration**

Our approach consists of four steps: (1) For each dataset, sample a representative subset of cells (atoms) that span both rare and abundant populations (2) For each dataset, learn a dictionary representation to reconstruct each cell, based on the atoms. (3) Integrate the atoms from each dataset (4) For each dataset, reconstruct each cell from the integrated atoms. Each step is described in detail below. We note that steps (1), (2), and (4) are performed on each dataset individually, and step (3) only requires performing joint computation on the downsampled set of atoms. Therefore, our procedure never requires loading or processing the entirety of the datasets at one time. Our approaches should therefore successfully extend to and beyond the analysis of 100,000,000 cells, which is now an achievable scale for combinatorial barcoding technologies.

All methods are implemented in our open-source R package Seurat (www.satijalab.org/seurat, www.github.com/satijalab/seurat)

1. *Sample a representative subset of cells ('atoms') from each dataset*

Our first step is to selectively downsample the cells in each dataset, aiming to identify a reduced set of cells that are representative of the full dataset. In particular, we aim to ensure that rare populations continue to be represented even after downsampling. We also aim to identify cell subsets in a computationally efficient manner, and to minimize any computation that must be performed on the full dataset prior to downsampling. We aim to select a subset of *k* cells from each dataset, each of which is referred to as an atom. In this manuscript we use *k*=5,000 unless otherwise noted.

We define:
$X \in \mathbb{R}^{n_{scRNA-seq} \times d_{genes}}$ : count matrix for scRNA-seq

$S \in \mathbb{R}^{k \times n_{scRNA-seq}}$ : Sampling matrix for the dataset. Each row is one-hot row vector matrix indicating which cells are selected, i.e. $s_{i,j} = 1$ if cell *i* is the *j*th cell to be selected. $i = 1, 2, \ldots, n_{cells}$, $j = 1, 2, \ldots, k$.

$SX \in \mathbb{R}^{k \times d_{genes}}$ : scRNA-seq matrix, after downsampling to the *k* cells selected. We also call this matrix A, as it represents the 'atoms' selected from the original dataset.

We can use a variety of techniques to define the sketching matrix S. These include geometric sketching techniques such as geosketch[9] or Hopper[10], or fast clustering procedures such as mini-batch kmeans[11] followed by cluster-informed downsampling.

In this work, we select cells based on their statistical leverage scores, a method for selecting influential data points in a dataset. In the context of linear regression, statistical leverage represents the influence of an individual data point in determining the best least-squares fit. In this context, cells with high leverage scores will tend to make the largest contribution to the gene covariance matrix, and therefore reflect the importance of the cell's profile. The exact statistical leverage score for a cell can be computed via an eigen decomposition of the X matrix, but this is computationally inefficient. As an alternative, Clarkson & Woodruff (2017) propose a randomized algorithm that efficiently computes a fast approximation of statistical leverage[3]. This algorithm is attractive for single-cell sequencing analysis as it is highly scalable and runs efficiently on sparse datasets. Briefly, the algorithm amounts to constructing a 'randomized' sketch of the input matrix based on the Johnson–Lindenstrauss lemma, and then computing the Euclidean norms of the rows of that sketch. The algorithm is fully described here [3], but we note the key mathematical steps below:

For the randomized sketching matrix, we use the sparse random CountSketch matrix C, which consists of 0, 1, and -1 elements, and is defined here[12].

$C \in \mathbb{R}^{c \times n_{scRNA-seq}}$: sparse randomized CountSketch matrix

We then perform a QR decomposition
$C\,X = [Q, R]$

We then apply a Fast Johnson-Lindenstrauss Transformation using a very sparse random projection matrix $\Pi$[13]. We calculate this matrix using the RandPro package[14] in R ('li' projection function).

$Z = X \times (R^{-1} \times \Pi)$

We can now calculate the leverage score for each cell, which are the Euclidean norms of the rows of the Z matrix. We can also calculate a sampling probability for selecting each cell $i$ as an atom, based on the leverage scores.

$l_i = ||Z[i,]||_2^2$: Leverage score for cell $i$.

$p_i = \frac{l_i}{\Sigma_{j=1}^{n} l_j}$ : probability of selecting cell $i$ as an atom,

Finally, we sample $k$ cells as atoms based on these probabilities. As described above, this procedure results in a downsampled dataset in which only the atoms remain, which we name A.

<u>2. Learn a dictionary representation to reconstruct each cell, based on the atoms</u>

We aim to learn reconstructions of $X$ based on the atoms of $A$, while minimizing the error between the original and reconstructed values. Specifically, we aim to identify the matrix D where:

$D \in \mathbb{R}^{n_{scRNA-seq} \times k}$: dictionary representation of scRNA-seq dataset such that:

$$\underset{D}{argmin}(\ ||DA - X||_F^2 + ||D||_F^2)$$

As described previously, this optimization problem is analogous to matrix regression, and has a closed-form solution for calculating $D$

$$D = XA^{\dagger}$$

Where † represents the pseudoinverse of the matrix.

3. Integrate the atoms from each dataset

Let $i = 1, 2, \ldots, n_{dataset}$ represent the datasets to be integrated, and let $A_i$ represent the matrix of atoms that result from downsampling dataset $i$. Our goal is to harmonize the set of matrices $[A_1, A_2, \ldots, A_{n_{dataset}}]$.

This can be performed with a wide variety of existing tools for the harmonization of single-cell datasets. For example, Seurat, Harmony, LIGER, scVI, Scanorama, fastMNN, scVI, and scArches, all learn a shared low-dimensional space that jointly represents the datasets, and aligns cells in a matched biological state together. Our goal is therefore to learn:

$$[A_1^*, A_2^*, \ldots, A_{n_{dataset}}^*]$$

Where $A_i^* \in \mathbb{R}^{n_{scRNA-seq} \times d_{RNA}}$: harmonized space for scRNA-seq dataset $i$

In this manuscript, we utilize our previously developed anchor-based workflow to integrate the datasets using reciprocal PCA, which is optimized for integration tasks with large numbers of samples and cells ('Fast integration using reciprocal PCA': https://satijalab.org/seurat/articles/integration_rpca.html). The integration procedure returns a low-dimensional space that jointly represents atoms from all datasets.

4. Reconstruct each cell from the integrated atoms.

The last step is performed individually for each dataset. Let $i = 1, 2, \ldots, n_{dataset}$ represent the datasets to be integrated, and let $X_i$ represent the full scRNA-seq count matrix representing dataset $i$.

We reconstruct integrated values for each cell in dataset $i$, using the previously computed dictionary representation for the dataset, along with the harmonized space $A_i^*$

$$X_i^* = D_i A_i^* = X_i (A_i^{\dagger} A_i^*)$$

The collection of matrices $[X_1^*, X_2^*, \ldots, X_{n_{dataset}}^*]$ now represents a low-dimensional space which jointly represents all cells from all datasets. Since these matrices are low-dimensional, each of them can be simultaneously loaded into memory. These representations can be used as input for common downstream analytical tasks including tSNE or UMAP visualization, graph-based clustering, and the identification of developmental trajectories.

### Preprocessing details for each dataset

*Adult mouse frontal cortex and hippocampus Paired-tag dataset:*

The datasets from Zhu (2021)[15] are generated with Paired-Tag, which performs simultaneous profiling of histone modifications and cellular transcriptomes, and contains a total of 64,849 nuclei. We extracted three datasets for the histone modifications H3K27ac, H3K4me1, and H3K27me3. We used the gene expression matrices as quantified in the original experiment. For each epigenetic modification, the original manuscript quantified read densities in 5k bins. These were aggregated into larger peaks using the CombineTiles function in Signac, and aggregated peaks less than 1MB in size were retained. We retained cells with total RNA counts between 500 and 10,000. We apply SCTransform to normalize the gene expression data and TF-IDF to normalize the histone modification data. We use PCA (dimensions 1:30) and TF-IDF (dimensions 2:30, excluding the first dimension as this is typically correlated with technical metrics in ATAC-seq or scCUT&Tag data) to reduce the dimensionality of the RNA and histone modification modalities, and construct the weighted nearest neighbor (WNN) graph.
Data acquisition source: GEO GSE152020

*Human frontal cortex snmC-seq data:*
This human frontal cortex dataset is a snmC-seq dataset from Luo (2017)[16] and contains 2,784 nuclei. We use the non-CG methylation 100kb bin count matrices as quantified in the original experiment We apply SCTransform[17] to normalize the gene expression data and log-normalization to normalize the methylation data. As this dataset is used as a query dataset in this manuscript, we do not perform unsupervised dimensionality reduction on the methylation data.
Data acquisition source: GEO GSE97179
https://brainome.ucsd.edu/annoj/brain_single_nuclei/

*Human frontal cortex snmC2T-seq data*
This human frontal cortex dataset is a snmC2T-seq data from Luo (2019)[18] and contains 4,357 nuclei. We use the non-CG methylation 100kb bin count matrices as quantified in the original experiment. We apply SCTransform to normalize the gene expression data and log-normalization to normalize the methylation data. We use PCA to reduce the dimensionality to 30 for both datasets and construct the weighted nearest neighbor (WNN) graph.
Data acquisition source: GEO GSE140493

*BMMC multiome*
We collected a total of ten 10x multiome datasets from the NeurIPS Multimodal Single-Cell Data Integration challenge website, representing 32,368 paired single-nucleus profiles of transcriptome and chromatin accesssibility. We retained cells with total RNA counts between 1,000 and 10,000 and total ATAC peak counts between 2,000 and 30,000. We apply SCTransform to normalize the gene expression data and TF-IDF to normalize ATAC peak counts. We use PCA (dimensions 1:40) and TF-IDF (dimensions 2:40) to reduce the dimensionality of each modality and construct the weighted nearest neighbor (WNN) graph.
Data acquisition source: https://openproblems.bio/neurips_docs/data/dataset/

*Human bone marrow mononuclear cells (BMMC) ATAC-seq*
This human bone marrow dataset is a snATAC-seq data from Granja[19] (2019). As the reads were originally mapped to hg19, we used cellranger-atac v2 to remap fastq files to hg38. In each cell, we quantified the same set of peaks that were detected in the BMMC multiome dataset. After removing low quality cells, 26,159 are retained with total ATAC peaks < 50,000 and > 2000. We apply TF-IDF to normalize the

ATAC-seq data. As this dataset is used as a query dataset in this manuscript, we do not perform unsupervised dimensionality reduction on the ATAC-seq data.
Data acquisition source: GEO GSE139369

*Human PBMC* CyTOF *dataset:*
This human PBMC CyTOF dataset was generated by the COVID-19 Multi-omics Blood Atlas COMBAT consortium, and consists of 7.11 million cells with a panel of 47 antibodies. We removed cells from sepsis patients, yielding a remainder of 5.17 million cells. We use the normalized expression matrices as quantified in the original study. As this dataset is used as a query dataset in this manuscript, we do not perform unsupervised dimensionality reduction on the protein data.
Data acquisition source: https://zenodo.org/record/5139561

*Azimuth reference:*
Azimuth scRNA-seq references for the human bone marrow (297,627 cells), and the human motor cortex (159,738 cells) were downloaded from the Human Biomolecular Atlas Project (HuBMAP) portal. The portal includes descriptions of each public data source used when compiling the reference dataset, as well as a link to a Github repository and Docker Hub to reproduce the construction of the reference.
Data acquisition: azimuth.hubmapconsortium.org

*Lung single cell-RNA datasets atlas:*
Nineteen datasets profiling human lung samples using scRNA-seq were downloaded from publicly available sources (links for each source dataset are provided in Supplementary Table 1). Low-quality cells were filtered using uniform QC thresholds: cells with RNA counts between 300 and 100,000, as well as mitochondrial read percentages below 20%, were retained. Normalization was performed using Log Normalization implemented in Seurat. We use PCA (dimensions 1:40) to reduce the dimensionality of each dataset.
Data acquisition source: Supplementary Table 1, lung scRNA datasets[20-38]

*PBMC COVID single-cell RNA datasets atlas:*

Fourteen datasets profiling human PBMC samples using scRNA-seq were downloaded from publicly available sources (links for each source dataset are provided in Supplementary Table 1). Eleven of these datasets had been previously organized in Tian (2022) [39]. Low-quality cells were filtered using uniform QC thresholds: cells with RNA counts between 150 and 150,000, as well as mitochondrial read percentages below 15%, were retained. Normalization was performed using Log Normalization implemented in Seurat. We use PCA (dimensions 1:40) to reduce the dimensionality of each dataset.
Data acquisition source: Supplementary Table 1, PBMC scRNA datasets[2,40-52]

### Differentiation trajectory and pseudotime analysis

In Fig. 2, we identify a myeloid differentiation trajectory and pseudotime ordering of cells that describes both reference (scRNA-seq) and query (scATAC-seq) cells. We extracted reference cells belonging to HSC, LMPP, GMP, and CD14 monocyte populations, and query cells that mapped to any of these subsets after bridge integration. We next constructed a *k*-nearest graph representing cells from both modalities, using the latent space learned during the bridge integration procedure. This graph was used as input to the destiny package, which reduces the dimensionality of the data using diffusion maps[53]. We note that as we manually selected cell populations that are known to encompass monocytic differentiation, we did not expect or observe branching events. We used the first two diffusion map coordinates as input to monocle3[54] in order to infer a pseudotemporal ordering.

We next aimed to identify cases where dynamic gene expression patterns ‘lag’ behind the accessibility dynamics of nearby regulatory regions. We can perform this analysis since our pseudotemporal ordering encompasses both scATAC-seq and scRNA-seq cells. We first associated each scATAC-seq peak with a gene using the ClosestFeature function in Signac. For each gene, we next smooth the expression profile along the learned trajectory using the ksmooth function (‘stats’ package in R[55]), using 1,000 intervals and a bandwidth of 0.01. We repeat the same process for the accessibility of each peak linked to this gene (bandwidth of 0.05). We next calculate the cross-correlation of the smoothed expression and accessibility values, which measures the similarity for the two time-series and calculates the optimal displacement of one relative to the other. We use the ccf function (‘stats’ package in R[55]), and identify a total of 574 gene-peak pairs with a cross-correlation > 0.6. Of these, we identify 236 cases exhibiting an optimal displacement of > 0.01 (we illustrate 6 such cases in Fig. 2l).

**Bridge cells down-sampling analysis**

To explore how the size and composition of the multi-omic dataset affected the robustness of bridge integration, we performed 25 serial downsamplings of the entire BMMC multi-omic dataset (200, 300, 400, 500, 600, 700, 800, 900, 1000, 2000, 3000, 4000, 5000, 6000, 7000, 8000, 9000, 10000, 11000, 12000, 13000, 14000, 15000, 20000, 30000). We used one batch of the scATAC dataset (12,256 cells) as a query, repeated bridge integration, and compared the resulting predictions with our original findings. As expected, we found that the degree of agreement after downsampling was cell-type dependent, as cells from abundant cell types were more robust to downsampling. We therefore expressed our results as a function of the number of cell types present in the bridge dataset for each cell type. For example, the 7,000 downsampled dataset contained 144 CD16 monocytes (Prediction accuracy: 1.00), and 22 pro-B cells (Prediction accuracy: 0.66). The 2,000 cell downsampled dataset contained 41 CD16 monocytes (Prediction accuracy: 0.94) and 6 pro-B cells (Prediction accuracy: 0.55). We aggregated all these results across downsamples and display the results in Fig 2a. For visual clarity, we only show an x-axis range of 10 to 500 in Fig. 2a.

**Benchmark analysis with multiVI and Cobolt**

In order to assess the performance of our bridge integration method alongside other recently proposed integration tools, we compared our results with multiVI [56] from scvi-tools v 0.14.5, and Cobolt[57] (v1.0.0). As both Cobolt and multiVI utilize variational autoencoders, both methods were run on a server with a discrete NVIDIA A100GPU with 40GB memory and pyTorch-lightning v.1.3.8 installed. Seurat analyses were run on a Intel Xeon Platinum 8280L server and used a single computational core.

For multiVI, we use the scRNA-seq, scATAC-seq and multi-omic RNA-ATAC paired counts matrices as input. We use the multiome_anndatas function to generate one anndata object for integration. We set batch information in categorical_covariate_keys, using the setup_anndata function. We then integrate the datasets by running the multiVI function, as outlined in the multiVI tutorial (https://docs.scvi-tools.org/en/stable/tutorials/notebooks/MultiVI_tutorial.html). We use 500 epochs for model training, as suggested in the multiVI tutorial. All other parameters were set to default settings. multiVI learns a latent space which jointly represents cells across the scRNA-seq and scATAC-seq datasets. We extract this space and perform nearest-neighbor calculations and UMAP visualization in Seurat.

For Cobolt, we use the scRNA-seq, scATAC-seq and multi-omic RNA-ATAC paired counts matrices as input. We use the SingleData function from cobolt_utils to generate three Cobolt objects and train the model using 20 latent variabes, a 0.001 learning rate, and 100 iterations as recommended in the Cobolt tutorial (https://github.com/epurdom/cobolt/blob/master/docs/tutorial.ipynb). All other parameters were set to default or recommended settings in the tutorial. Cobolt learns a latent space which jointly represents cells across the scRNA-seq and scATAC-seq datasets. We extract this space and perform nearest-neighbor calculations and UMAP visualization in Seurat.

We performed comparative benchmarking in three contexts. First, we ran all three approaches on the datasets from Fig. 2, aiming to map an scATAC-seq query dataset onto an scRNA-seq defined reference. We do not have ground truth information for this dataset, so we do not calculate quantitative benchmarks, though we visualize the performance of all methods in Fig. 3b and Supplementary Fig. 2. As multiVI and Cobolt do not provide methods to explicitly label query scATAC-seq cells using scRNA-seq references, we employed a commonly used heuristic for label transfer: for each scATAC-seq cell we identified the closest five neighbors in scRNA-seq cells, and transferred the most common cell annotation amongst the neighbors. In Fig. 3b, we visualize chromatin accessibility at the SIGLEC6 locus for cells predicted as ASDC by all methods, and we show additional loci in Supplementary Fig. 2.

Second, we performed quantitative benchmarking in a context where we had ground-truth dataset to establish the accuracy of scATAC-seq / scRNA-seq integration. We split the BMMC multi-omic dataset into two groups. The first group consists of a randomly sampled subset of 2,115 cells representing at most 100 cells per author-defined celltype. This group of cells was utilized as the multi-omic bridge dataset for benchmarking. The remaining cells were placed in the second group, and were split into separate scRNA-seq and scATAC-seq profiles (i.e. the multi-omic pairing information was temporarily discarded). We then integrated the datasets using either bridge integration (using both Seurat v3 and mnnCorrect for the final alignment step), multiVI, or Cobolt. After integration, all methods return a latent space that jointly represents cells from both the scATAC-seq and scRNA-seq datasets. For each scATAC-seq cell, we know its matched scRNA-seq profile, as they were originally measured simultaneously. Successful integration techniques will place matched profiles place close together in this latent space. For each scATAC-seq cell, we therefore calculated the Jaccard similarity metric to its matched scRNA-seq profile (we note that this similarity metric is symmetric). We report these results in Fig. 3c and Supplementary Fig. 2, either averaged together across all cells, or averaged within author-defined cell types.

Third, we repeat the ground-truth benchmarking analysis on a second multi-omic technology. Paired-tag enables simultaneous CUT&Tag and transcriptomic profiling in single cells. We used data for three histone modifications: H3K27ac, H3K27me3, and H3K4me1. As each dataset consists of multiple replicates, we used replicate 1 as the multi-omic dataset, and split the CUT&Tag and RNA modalities from the second replicate for benchmarking. We ran multiVI, Cobolt, and bridge integration (using both Seurat v3 and mnnCorrect for the final alignment step) as before, substituting the CUT&Tag counts matrix for the scATAC-seq matrix as previously described.

**Community-wide integration analyses**

To facilitate the harmonization and subsequent meta-analysis of a diversity of publicly available scRNA-seq datasets, we apply our atomic sketch integration approach to 1,525,710 scRNA-seq profiles spanning 19 publicly available human lung scRNA-seq datasets. As described above, we calculate a leverage score for each cell in each dataset, and use this to sample 5,000 cells as atoms. We find that these 5,000 cells retain rare cell types, despite downsampling (Supplementary Fig. 3). We learn a dictionary representation that reconstructs cells from each dataset based on the selected atoms using the methods described above. We use our previously developed reciprocal PCA-based integration workflow (https://satijalab.org/seurat/articles/integration_rpca.html) to integrate the 95,000 atoms originating from these 19 datasets. Finally, the learned dictionary representations can be used to reconstruct harmonized profiles (in low-dimensional space) for all 1,525,710 scRNA-seq profiles. This space was used as input for UMAP to generate the visualization in Fig. 4b,c.

The harmonized space for all 1,525,710 scRNA-seq can also be used as input to graph-based clustering approaches. However, since annotation is an iterative and manual process, we chose to first perform clustering on the harmonized dataset of 95,000 atoms. We constructed a shared nearest-neighbor graph, and partition this into clusters using the graph-based smart local moving (SLM) algorithm[58]. We initially clustered cells at a high resolution (resolution = 5) and performed differential expression on all pairs of clusters for RNA markers. We merged clusters that did not exhibit clear evidence of separation. We removed clusters that showed clear evidence of expressing markers for two different cell types as likely doublets. To assign names to individual clusters, we used the recently published Anatomical Structures, Cell Types, plus Biomarkers (ASCT+B) tables[59], except for five clusters (Adventitial Fibroblast, Alveolar Fibroblast, Myofibroblast, proliferating NK/T, Squamous), where our desired annotation was not present in the most recently available version of the table (v1.1[60]). For each cell in the full dataset, we find its 10 nearest neighbors amongst the annotated atoms, and transfer the most commonly observed annotation.

In Fig. 5, we perform 'community-wide' integration on 3.46M cells spanning 639 individuals and 14 studies. As these studies varied widely in the number of cells present in each dataset, we selected at least 5,000 and at most 10% of the cells in each dataset as atoms based on their leverage score. This enabled the larger and more comprehensive datasets to contribute additional weight to the integrated reference. We performed integration, reconstruction, and annotation using the same steps as described for the lung.

**Identifying differentially expressed genes across cell types and conditions.**

In the lung and PBMC community-wide integration, we identify differentially expressed (DE) genes on the 'pseudobulk' expression values calculated from each of individual study. We perform logistic regression-based method to identify DE genes. For space considerations, we typically report only the top 10 markers in each heatmap, and sort genes first by adjusted p-value and next by log fold-change to determine the top markers. In order to compare the results of single-cell and bulk analyses, we used the wilcoxauc method from presto[61] to identify DE genes using either the single-cell or pseudobulk profiles as input, and sorted by the AUC statistic. In Fig. 4g, we compare the distribution of average expression values (within a cell-type) for the top 100 markers identified by either single-cell or pseudobulk analysis.

To identify COVID-19 response signatures that are consistent across multiple patients, we first calculate a pseudobulk average for CD14 monocytes for each of the 506 donors who are either healthy controls, or whose metadata indicated mild, moderate, or severe COVID-19[39]. We performed DE analysis at the pseudobulk level to identify markers of CD14 monocytes expressed in severe COVID samples compared to healthy controls. In Fig. 5b, we order each pseudobulk profile by the expression level of these genes, which are enriched for interferon response genes, for visualization. We repeat this process for 8 additional cell states in Supplementary Fig 5b.

### REFEERENCES


1. Barshan, E., Ghodsi, A., Azimifar, Z. & Jahromi, M.Z. Supervised principal component analysis: Visualization, classification and regression on subspaces and submanifolds. *Pattern Recognition* **44**, 1357-1371 (2011).
2. Hao, Y. et al. Integrated analysis of multimodal single-cell data. *Cell* (2021).
3. Clarkson, K.L. & Woodruff, D.P. Low-rank approximation and regression in input sparsity time. *Journal of the ACM (JACM)* **63**, 1-45 (2017).
4. Woodruff, D.P. Sketching as a tool for numerical linear algebra. *arXiv preprint arXiv:1411.4357* (2014).
5. Belkin, M. & Niyogi, P. Laplacian eigenmaps for dimensionality reduction and data representation. *Neural computation* **15**, 1373-1396 (2003).
6. Levine, J.H. et al. Data-Driven Phenotypic Dissection of AML Reveals Progenitor-like Cells that Correlate with Prognosis. *Cell* **162**, 184-197 (2015).

34. Consortium, T.T.S. & Quake, S.R. The Tabula Sapiens: a multiple organ single cell transcriptomic atlas of humans. *bioRxiv*, 2021.2007.2019.452956 (2021).
35. Travaglini, K.J. et al. A molecular cell atlas of the human lung from single-cell RNA sequencing. *Nature* **587**, 619-625 (2020).
36. Wang, A. et al. Single-cell multiomic profiling of human lungs reveals cell-type-specific and age-dynamic control of SARS-CoV2 host genes. *Elife* **9** (2020).
37. Watanabe, N. et al. Anomalous epithelial variations and ectopic inflammatory response in chronic obstructive pulmonary disease. *medRxiv*, 2020.2012.2003.20242412 (2021).
38. Wauters, E. et al. Discriminating mild from critical COVID-19 by innate and adaptive immune single-cell profiling of bronchoalveolar lavages. *Cell Res* **31**, 272-290 (2021).
39. Tian, Y. et al. Single-cell immunology of SARS-CoV-2 infection. *Nat Biotechnol* **40**, 30-41 (2022).
40. Arunachalam, P.S. et al. Systems biological assessment of immunity to mild versus severe COVID-19 infection in humans. *Science* **369**, 1210-1220 (2020).
41. Ahern, D.J. et al. A blood atlas of COVID-19 defines hallmarks of disease severity and specificity. *MedRxiv* (2021).
42. Combes, A.J. et al. Global absence and targeting of protective immune states in severe COVID-19. *Nature* **591**, 124-130 (2021).
43. Lee, J.S. et al. Immunophenotyping of COVID-19 and influenza highlights the role of type I interferons in development of severe COVID-19. *Sci Immunol* **5** (2020).
44. Ren, X. et al. COVID-19 immune features revealed by a large-scale single-cell transcriptome atlas. *Cell* **184**, 5838 (2021).
45. Schulte-Schrepping, J. et al. Severe COVID-19 Is Marked by a Dysregulated Myeloid Cell Compartment. *Cell* **182**, 1419-1440 e1423 (2020).
46. Silvin, A. et al. Elevated Calprotectin and Abnormal Myeloid Cell Subsets Discriminate Severe from Mild COVID-19. *Cell* **182**, 1401-1418 e1418 (2020).
47. Stephenson, E. et al. Single-cell multi-omics analysis of the immune response in COVID-19. *Nat Med* **27**, 904-916 (2021).
48. Su, Y. et al. Multi-Omics Resolves a Sharp Disease-State Shift between Mild and Moderate COVID-19. *Cell* **183**, 1479-1495 e1420 (2020).
49. Wilk, A.J. et al. A single-cell atlas of the peripheral immune response in patients with severe COVID-19. *Nat Med* **26**, 1070-1076 (2020).
50. Yao, C. et al. Cell-type-specific immune dysregulation in severely ill COVID-19 patients. *Cell Rep* **34**, 108943 (2021).
51. Yu, K. et al. Dysregulated adaptive immune response contributes to severe COVID-19. *Cell Res* **30**, 814-816 (2020).
52. Zhu, L. et al. Single-Cell Sequencing of Peripheral Mononuclear Cells Reveals Distinct Immune Response Landscapes of COVID-19 and Influenza Patients. *Immunity* **53**, 685-696 e683 (2020).
53. Haghverdi, L., Buettner, F. & Theis, F.J. Diffusion maps for high-dimensional single-cell analysis of differentiation data. *Bioinformatics* **31**, 2989-2998 (2015).
54. Qiu, X. et al. Reversed graph embedding resolves complex single-cell trajectories. *Nat Methods* **14**, 979-982 (2017).
55. Team, R.C. (Vienna, Austria, 2013).
56. Ashuach, T., Gabitto, M.I., Jordan, M.I. & Yosef, N. MultiVI: deep generative model for the integration of multi-modal data. *bioRxiv*, 2021.2008.2020.457057 (2021).
57. Gong, B., Zhou, Y. & Purdom, E. Cobolt: integrative analysis of multimodal single-cell sequencing data. *Genome Biol* **22**, 351 (2021).
58. Waltman, L. & Van Eck, N.J. A smart local moving algorithm for large-scale modularity-based community detection. *The European physical journal B* **86**, 471 (2013).
59. Borner, K. et al. Anatomical structures, cell types and biomarkers of the Human Reference Atlas. *Nat Cell Biol* **23**, 1117-1128 (2021).
60. Gloria Pryhuber, X.S. (2021).
61. Korsunsky, I., Nathan, A., Millard, N. & Raychaudhuri, S. Presto scales Wilcoxon and auROC analyses to millions of observations. *BioRxiv*, 653253 (2019).
